# TET3 protects the Dlk1-Dio3 Imprinted Locus from DNA hypomethylation during adult NSC Reprogramming

**DOI:** 10.1101/2025.02.13.638077

**Authors:** Laura Lázaro-Carot, Esteban Jiménez-Villalba, Jordi Planells, Anna Lozano-Ureña, Jennifer Díaz-Moncho, Raquel Montalbán-Loro, Adela Lleches-Padilla, Martina Kirstein, Mitsuteru Ito, Elizabeth J. Radford, Sacri R. Ferrón

## Abstract

Genomic imprinting is an epigenetic mechanism that drives monoallelic gene expression depending on parental origin. Loss of imprinting (LOI) is associated with human imprinting disorders, fetal development, and cancer progression. Imprinted genes, organized in clusters, are regulated by methylation at imprint control regions (ICRs), differentially methylated regions (DMRs) between parental chromosomes. Somatic cell reprogramming into induced pluripotent stem cells (iPSCs) is a valuable tool for studying pluripotency and holds promise for patient-specific therapies. Discerning whether genomic imprinting changes during reprogramming represent epigenetic abnormalities or essential adaptations linked to pluripotency is crucial. Here, we perform RNA-seq and MeDIP-seq analysis on mouse iPSCs derived from neural stem cells (NSCs). Our findings reveal that ICRs undergo DNA hypomethylation, confirming widespread LOI in pluripotent cells. However, the IG-DMR within the Dlk1-Dio3 imprinted cluster resists hypomethylation, a hallmark of successful pluripotency acquisition. We also identify a non-canonical role of TET3 in IG-DMR methylation protection through transcriptional regulation of *Oct4* and *Trim28*. These findings highlight genomic imprinting as a key mechanism of gene dosage control in pluripotency acquisition and maintenance.

## Introduction

During mammalian development, the vast majority of genes are expressed or repressed from both alleles. However, there is a small number of genes, termed “*imprinted genes*” that are expressed monoallelicaly from either the maternally or the paternally inherited chromosomes (Ferguson-Smith, 2011). Approximately 150 imprinted genes have been described in mammals and are generally organized in clusters, although examples of singleton imprinted genes do exist (Barlow & Bartolomei, 2014; Lassi & Tucci, 2019; Ferguson-Smith, 2011). An imprinting cluster is usually under the control of a DNA element, called the imprinting control region (ICR) that consists of differentially DNA methylated regions (DMRs) between the two parental chromosomes (Edwards & Ferguson-Smith, 2007; SanMiguel & Bartolomei, 2018; Tucci *et al*, 2019). Deletion or alteration of an ICR results in loss of imprinting (LOI) of multiple genes in the cluster and this has been associated to several human pathologies and imprinting disorders such as Prader Willi Syndrome (PWS) and Angelman Syndrome (Barlow & Bartolomei, 2014; Ferguson-Smith, 2011; Lassi & Tucci, 2019). The parental specific marks at ICRs are established in the developing germline, giving as a result gametes bearing imprints according to the sex of the individual (Bartolomei & Ferguson-Smith, 2011; Edwards & Ferguson-Smith, 2007; Ferguson-Smith, 2011). After fertilization, a rapid and extensive reprogramming of the parentally inherited genomes occurs, and most DNA methylation is lost (Smallwood & Kelsey, 2012; SanMiguel & Bartolomei, 2018). However, the parental-specific imprints are maintained during this period and a memory of parental origin is propagated into daughter cells during somatic cell divisions (Takahashi *et al*, 2015). At a molecular level, DNA methylation is a dynamic process where different enzymes are involved in actively methylating and demethylating cytosine residues of the DNA. Methylation is established and maintained by DNA methyltransferase (DNMT) family of enzymes, either *de novo* by DNMT3A and DNMT3B, or maintained by DNMT1. On the other hand, ten-eleven translocases (TET) enzymes family, integrated by TET1, TET2 and TET3, catalyse the conversion of 5-methylcytosine (5mC) into 5-hydroxymethylcytosine (5hmC) to remove methylation marks (Wu & Zhang, 2017). TET proteins have been implicated in maintaining DNA methylation at ICRs during the germline resetting of genomic imprints in embryonic development (Hackett *et al*, 2013). In addition to their well-characterized catalytic function, TET proteins bind to 5mC-free promoters and interact with key epigenetic regulators, including histone deacetylases, acetyltransferases and Polycomb repressive complex 2, to regulate gene expression (Lian *et al*, 2016). This suggests additional functions for TET enzymes independent of their catalytic activity (Montalbán-Loro *et al*, 2019).

*In vitro* reprograming of somatic cells into induced pluripotent stem cells (iPSCs), has enormous therapeutic potential as it opened up the possibility of generating patient-specific pluripotent cell lines to study and treat different degenerative diseases. A variety of cell types have been reprogrammed into iPSCs including fibroblasts (Takahashi & Yamanaka, 2006), hepatocytes, gastric epithelial cells, B cells (Hanna *et al*, 2008), pancreatic β cells (Stadtfeld *et al*, 2008), neural progenitor cells (Kim *et al*, 2008a), melanocytes or keratinocytes (Aasen *et al*, 2008). Different combinations of reprogramming factors, such as *Oct4*, *Sox2*, *Klf4* and *c-Myc*, have been used to convert somatic cells into iPSCs with comparable expression profiles to embryonic stem cells (ESCs) (Eminli *et al*, 2008; Kim *et al*, 2008a). The key determinants and the temporal sequence of epigenetic events that transition a differentiated cell to a pluripotent state during iPSC derivation remain incompletely understood.

Evidence from multiple studies highlights the critical role of DNA methylation changes in successful reprogramming, particularly the necessity for demethylation at promoters of pluripotency-associated genes (Takahashi & Yamanaka, 2006). Therefore, incomplete DNA demethylation can result in only partially reprogrammed cells (Mikkelsen *et al*, 2008), as the erasure of differentiation-specific epigenetic marks is required for faithful reprogramming (Apostolou *et al*, 2013). While genomic imprinting remains relatively stable in somatic cells, it is variably lost during iPSCs reprogramming, with some imprinted regions more severely affected than others (Arez *et al*, 2022; Kim *et al*, 2013; Lee *et al*, 2016; Liu *et al*, 2010; Yagi *et al*, 2019; Takikawa *et al*, 2013; Perrera & Martello, 2019). Moreover, aberrant silencing of imprinted genes during iPSC generation has been linked to impaired tissue development (Yagi *et al*, 2019; Li *et al*, 2019). Several studies have reported global hypomethylation of ICRs during reprogramming, although *de novo* methylation at later stages has also been observed (Yagi *et al*, 2019). For instance, the *Dlk1-Dio3* imprinted region on murine chromosome 12 has been shown to undergo hypermethylation in iPSCs, disrupting the expression of multiple genes within the cluster (Stadtfeld *et al*, 2008; Liu *et al*, 2010; Stadtfeld *et al*, 2010; Pham *et al*, 2022). Additionally, imprinting defects in iPSCs appear to depend on both the sex of donor cells and culture conditions, and these defects cannot be rescued upon differentiation (Arez *et al*, 2022; Nazor *et al*, 2012). Most of these studies have analysed only a limited number of imprinted genes and ICRs in iPSCs, providing a partial view of genomic imprinting alterations. Therefore, distinguishing between essential and undesirable imprinting changes in iPSCs is crucial for their therapeutic applications.

Here, we present a comprehensive and unbiased analysis of transcriptome and methylome alterations in iPSCs derived from adult murine NSCs, using only the two reprogramming factors *Klf4* and *Oct4*. Genome-wide RNA-seq and MeDIP-seq analysis reveal a profoundly altered transcriptome in iPSCs, accompanied by extensive DNA hypomethylation. Studying DNA methylation across all described ICRs shows that most DMRs undergo hypomethylation, confirming a global loss of genomic imprinting in the iPSCs generated from adult NSCs. These methylation changes strongly correlate with transcriptional alterations in genes within affected imprinted clusters. However, the IG-DMR, which regulates the *Dlk1-Dio3* imprinting cluster on chromosome 12, resists this widespread hypomethylation during reprogramming, preserving genomic imprinting at this locus in iPSCs. We propose a model in which TET3 transcriptionally regulates the expression of *Trim28* and *Oct4* to safeguard IG-DMR methylation.

## Results

### NSCs from the adult SVZ convert into a pluripotent state only with the transduction of *Oct4* and *Klf4*

Previous studies have reported that neurosphere cultures obtained from postnatal day 5 mouse brain endogenously express *Sox2*, *c-Myc* and *Klf4*. Thus, these cultures can be reprogrammed with *Oct4* alone, or with *Oct4* and *Klf4* at a similar efficiency to the reprogramming rate of murine fibroblast with the original four factors (Kim *et al*, 2008b). To assess the expression level of these transcription factors in NSCs derived from the adult subventricular zone (SVZ), quantitative PCR (qPCR) was performed using a cell line of pluripotent embryonic stem cells (ESCs) as a reference. We found that adult NSCs consistently expressed neural genes such as *Pax6* or *Olig2*, and also expressed high levels of *Sox2*, *Klf4*, and *c-Myc* (**Fig. 1A**). As expected, genes associated with pluripotency, such as *Oct4*, *Nanog* or *Zfp42* were not expressed in adult NSCs (**Fig. 1A**). Based on this gene expression profile, we developed a reprogramming protocol using retroviral vectors encoding only the transcription factors *Oct4* and *Klf4* (2 factors condition, 2F), along with a retrovirus encoding the red fluorescent protein *mCherry* to track the exogenous expression of the reprogramming factors (**Fig. 1B**). These retroviruses were produced by transfecting Plat-E packing cells with the retroviral plasmids (**Fig. 1B** and **S1A**). Post-infected NSCs (PI-NSCs) were grown on a feeder layer of mouse embryonic fibroblasts using a medium supplemented with the cytokine leukaemia inhibitory factor (LIF) (**Fig. 1B**). After 10 days, cultures started to form clone-like aggregates, which were large, with poorly defined edges and some of them expressed the pluripotency marker stage-specific embryonic antigen 1 (SSEA1) (**Fig. 1B** and **S1A**). Retroviral vectors are transcriptionally silent in pluripotent stem cells (Hotta & Ellis, 2008). However, most of the clones formed were still positive for mCherry, indicating that despite expressing SSEA1, they were only partially reprogrammed (**Fig. S1A**). We considered this state of cells as pre-iPSCs (**Fig. 1B** and **S1A**). To promote a ground state of pluripotency of these pre-iPSCs, we applied molecularly defined conditions by neutralizing inductive differentiation stimuli with a dual inhibition (2i) of mitogen-activated protein kinase signalling (MEK) and glycogen synthase kinase-3 (GSK3) (Silva *et al*, 2008). In this serum-free culture medium, LIF was also added, to maximize clonogenic self-renewal of pluripotent cells (Ying *et al*, 2008) (**Fig. 1B)**. This reprogramming protocol was repeated in 6 independent cultures of adult NSCs. After 10 days in the new controlled 2i/LIF conditions, ESC-like colonies that did not express mCherry were obtained (**Fig. S1B**), suggesting that cells had been fully reprogrammed into iPSCs. At least 10 clones of each culture were isolated and expanded i*n vitro* (**Fig. 1B).**

**FIGURE 1.**
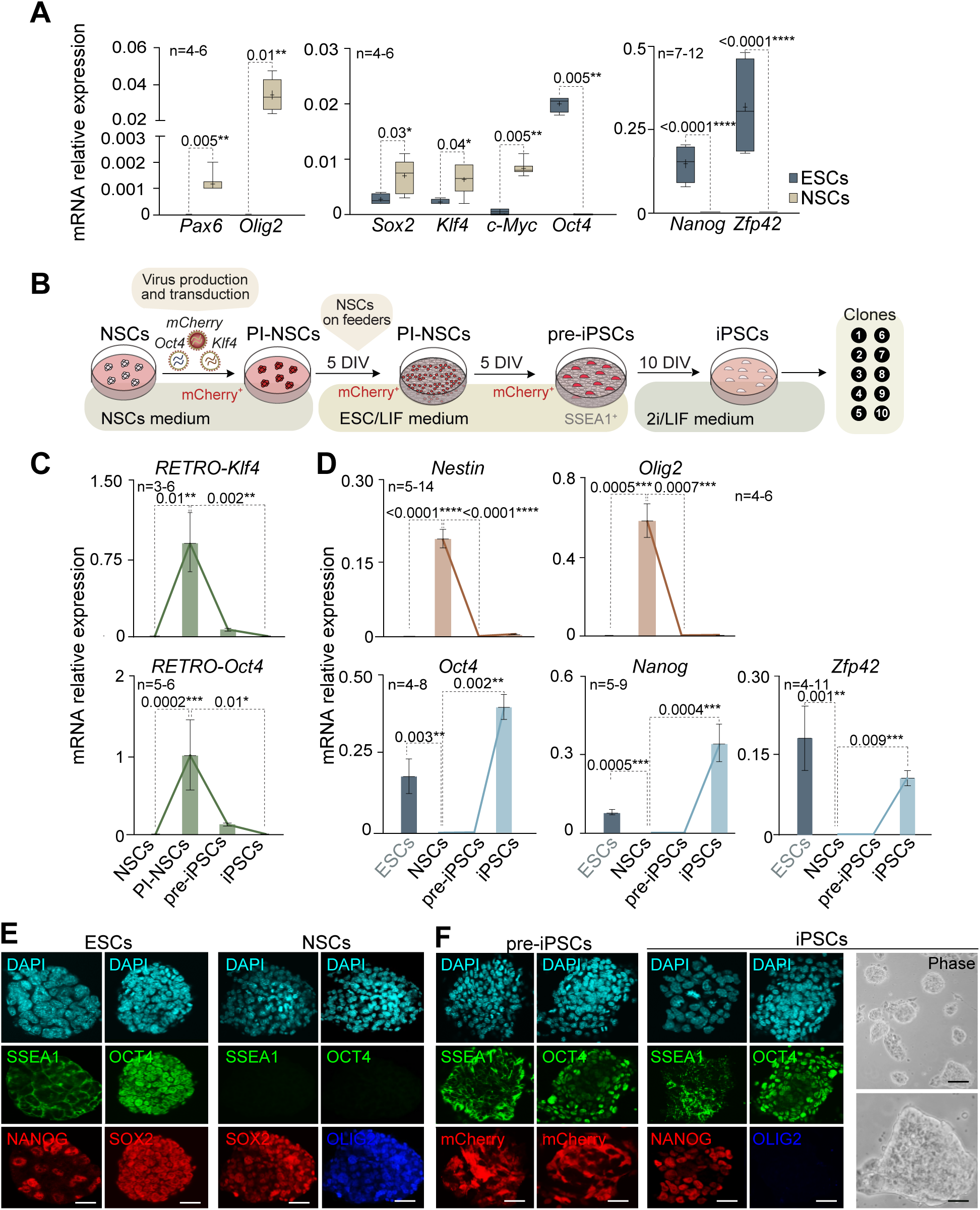
NSCs from the adult SVZ are reprogrammed into iPSCs by exogenous expression of *Oct4* and *Klf4*. **(A)** Quantitative PCR (qPCR) of the neural genes *Pax6* and *Olig2,* and the pluripotency genes *Nanog* and *Zfp42* in ESCs (blue) and adult NSCs (beige) (left and right panels). qPCR analysis of the endogenous expression of the reprogramming transcription factors *Oct4*, *Sox2*, *Klf4* and *c-Myc* in ESCs and adult NSCs is also shown (middle panel). **(B)** Schematic representation of the protocol used to reprogram adult NSCs into iPSCs. NSCs are infected with retroviruses encoding *Oct4*, *Klf4* and the fluorescent protein mCherry. After 5 days *in vitro* (DIV) in NSCs medium, neurospheres formed by post-infected NSCs (PI-NSCs) are dissociated into single cells and plated on murine embryonic fibroblasts using ESC/LIF medium. 5 days after dissociation, mCherry^+^ and SSEA1^+^ clone-like aggregates containing pre-iPSCs start to appear. Medium is then changed to 2i/LIF medium to complete the reprogramming process. After 10 more DIVs, cells have become full iPSCs and 10 single clones of each culture are picked and subcultured independently for further analysis. **(C)** qPCR expression analysis of retroviral *Klf4* and *Oct4* expression in adult NSCs, PI-NSCs, pre-iPSCs and iPSCs**. (D)** qPCR expression analysis of the neural genes *Nestin* and *Olig2* (upper panel) and the pluripotency-related genes *Oct4*, *Nanog and Zfp42* (lower panel) in NSCs, pre-iPSCs and iPSCs. ESCs were used as a control of the pluripotent state. **(E)** Immunocytochemistry (ICC) images of SSEA1, OCT4 (green) and SOX2 (red) in ESCs (left panel) and adult NSCs (right panel). ICC for the pluripotency marker NANOG (red) for ESCs (left panel) and for the neural marker OLIG2 (blue) in NSCs (right panel) are also shown. **(F)** ICC images for SSEA1 and OCT4 (green) in pre-iPSCs (left panel) and iPSCs (middle panel). mCherry fluorescence for pre-iPSCs (left panel) and ICC for the pluripotency marker NANOG (red) and the neural marker OLIG2 (blue) for iPSCs (middle panel) are shown. Phase contrast images for fully reprogrammed iPSCs clones (right panel) are also shown. *Gapdh* was used as a housekeeping gene for qPCR normalization. DAPI was used to counterstain nuclei in immunofluorescence images. Scale bars for ICCs in e and f: 20 μm; and for phase contrast images in e 40 μm in upper panel and 5 μm in lower panel. P-values and number of samples are indicated. In box and whiskers plots, the mean is indicated as + and whiskers represent the maximum and minimum values. In bar plots, error bars show s.e.m.

To further characterise the acquisition of a pluripotent state of these clones, we performed a qPCR analysis on the PI-NSCs, pre-iPSCs and the iPSCs generated. As expected, PI-NSCs expressed high levels of both *Oct4* and *Klf4* retroviral transgenes (**Fig. 1C**). Although lower, retroviral genes expression was still present in pre-iPSCs (**Fig. 1C**). The expression levels of the neural markers *Nestin* and *Olig2* were downregulated in pre-iPSCs, although the expression of pluripotency markers such *Oct4*, *Nanog* and *Zfp42* remained practically undetectable in these cells (**Fig. 1D**). These results confirmed that pre-iPSCs were an intermediate state in which critical attributes of true pluripotency, including stable expression of endogenous *Oct4* and *Nanog,* were not attained yet. Complete downregulation of retroviral transgenes, essential for full reprogramming, was corroborated in finally derived iPSCs compared to the original infected NSCs and to pre-iPSCs (**Fig. 1C**). This was accompanied by the stable induction of endogenous pluripotency-related genes *Oct4, Nanog* and *Zfp42* (**Fig. 1D**) consistent with the acquisition of a pluripotent state. Moreover, the expression of neural specific genes such as *Nestin* and *Olig2*, was completely absent in iPSCs (**Fig. 1D**). Immunocytochemistry studies confirmed the presence of mCherry in pre-iPSCs but not in iPSCs (**Fig. 1E,F** and **S1B**). Immunocytochemistry for OCT4 and NANOG, together with histochemistry against alkaline phosphatase activity confirmed the pluripotency of the generated iPSCs (**Fig. 1F** and **S1C**).

We next corroborated the naive pluripotency of iPSCs by evaluating the reactivation of the inactive X chromosome in female iPSCs (Janiszewski *et al*, 2019). RNA levels of *Xist*, responsible for X chromosome inactivation were significantly reduced in iPSCs together with an increase in *Tsix* expression (**Fig. S1D**) confirming the X chromosome reactivation and thus the acquisition of a fully pluripotent state in iPSCs. Moreover, genetic variations, such as aneuploidy or polyploidy, may be introduced during the generation of iPSCs (Vaz *et al*, 2021). Karyotype analysis in iPSCs clones revealed that the majority of the analysed lines (93%) exhibited a normal karyotype with around 40 chromosomes per metaphase (**Fig. S1E**) and iPSCs lines with chromosomal abnormalities were excluded from further studies.

### iPSCs derived from adult NSCs differentiate *in vitro* and *in vivo* into cells from the three germ layers

Pluripotent stem cells have the potential to differentiate into cells of the three germ layers: mesoderm, endoderm and ectoderm. This differentiation potential is typically confirmed by demonstrating the capacity of the iPSCs to form three-dimensional structures called embryoid bodies (EBs), comprising characteristic cell types of these germ layers (Höpfl *et al*, 2004). EBs emulate the structure of developing embryos and serve as a model to obtain various cell lineages (Spelke *et al*, 2010). Thus, to explore the developmental capacity of the iPSCs generated from adult NSCs, we induced EBs formation by subjecting cells to conditions that are adverse to pluripotency and proliferation, using the hanging drop method (Höpfl *et al*, 2004) (**Fig. 2A**). Suspended iPSCs on the dish lid aggregate at the base of the drop consistently generating uniform EBs (**Fig. 2A**). ESCs and ESCs-derived EBs were utilized as a comparative condition of differentiation (**Fig. S2**).

**FIGURE 2.**
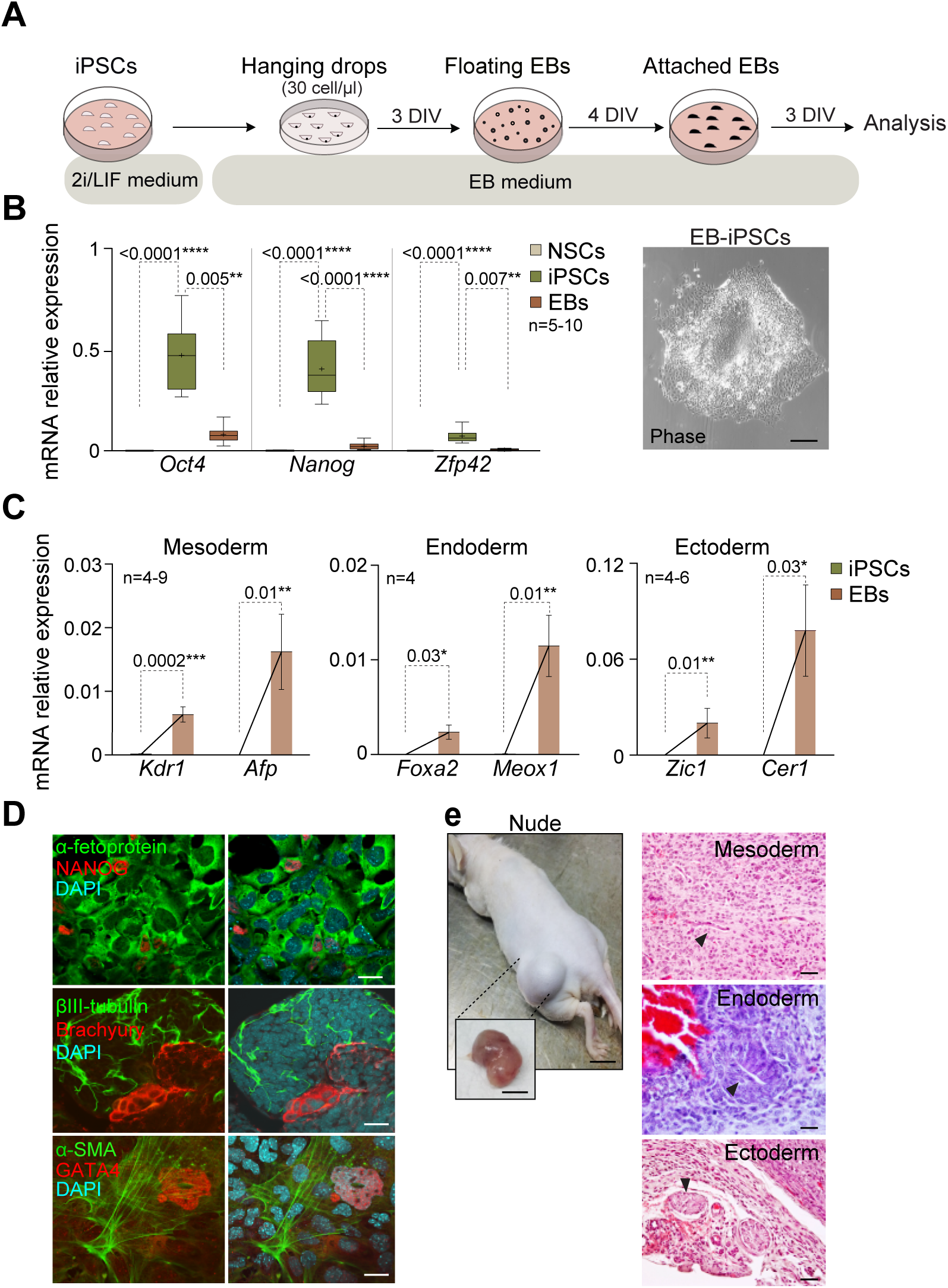
iPSCs generated from NSCs are able to differentiate into cells of the three germ layers *in vitro* and *in vivo*. **(A)** Schematic representation of the embryoid bodies (EB) assay using the “*hanging drops*” method. iPSCs were dissociated and the cell suspension (30 cells/μL) was distributed in drops in a plate that was incubated upside-down for 3 days *in vitro* (DIVs) in EB medium. Then, the plate was inverted and culture media was added. Incipient EBs were incubated 4 more DIVs in floating conditions. Then, EBs were seeded in gelatine pre-treated plates to allow differentiation for 3 more DIVs before analysis. **(B)** qPCR expression analysis of pluripotency-related genes *Oct4*, *Nanog* and *Zfp42* genes in NSCs (beige), iPSCs (green) and iPSCs-derived EBs (brown) (left panel). Phase contrast image of an EB is included (right panel). **(C)** qPCR analysis of *Kdr1* and *Afp* (mesoderm), *Foxa2* and *Meox1* (endoderm), and *Zic1* and *Cer1* (ectoderm) in iPSCs (green) and iPSCs-derived EBs (brown). **(D)** ICC detection of the pluripotency marker NANOG (red) and the different germ layer markers α-fetoprotein (green, endoderm) (upper panel), βIII-tubulin (green, ectoderm) and Brachyury (red, mesoderm) (middle panel), and α-SMA (green, mesoderm) and GATA4 (red, endoderm) (lower panel) in iPSCs-derived EBs. **(E)** Image of the dorsolateral area of immunocompromised *Nude* mice 2 weeks after the injection of iPSCs, including a detailed image of the formed teratoma after its extraction (left panel). Histological analysis of teratomas using haematoxylin-eosin staining (right panel). Muscle fibres derived from mesoderm, columnar epithelium derived from endoderm and epithelial cells derived from ectoderm are indicated with arrowheads. *Gapdh* was used as a housekeeping gene for qPCR normalization. DAPI was used to counterstain nuclei in immunofluorescence images. Scale bars in b: 10 μm; in d: 50 μm; and in e: 1 cm (left panel) and 20 μm (right panel). P-values and number of samples are indicated. In box and whiskers plots, the mean is indicated as + and whiskers represent the maximum and minimum values. In bar plots, error bars show s.e.m.

Initially, we determined the gene expression levels of pluripotency and differentiation genes in iPSCs-derived EBs using qPCR analysis. The results showed a downregulation for the pluripotency markers *Oct4*, *Nanog* and *Zfp42* upon differentiation (**Fig. 2B**). Additionally, a significant increase for the mesoderm markers *Kdr1* and *Afp*, the endoderm markers *Foxa2* and *Meox1,* and the ectoderm markers *Zic1* and *Cer1* was observed, confirming the presence of cells from the three germ layers within the generated EBs (**Fig. 2C** and **S2A)**. The presence of the differentiated cells from all three germ layers was further confirmed by immunocytochemistry with specific antibodies against α-fetoprotein and GATA4 (endoderm), βIII-tubulin (neuroectoderm), and Brachyury and α-SMA (mesoderm) (**Fig. 2D** and **S2B**). We next evaluated the *in vivo* pluripotent capacity of iPSCs derived from adult NSCs using the teratoma assay (Prokhorova *et al*, 2009). Teratomas are non-malignant tumours that result from uncontrolled expansion and disorganized differentiation of pluripotent cells. When ESCs are transplanted into immunocompromised *Nude* mice, they trigger teratoma formation (Prokhorova *et al*, 2009). In this study, adult NSCs-derived iPSCs were injected into the dorsolateral area in the subcutaneous space of *Nude* mice (**Fig. 2E**). Subsequent histological analysis of the teratomas, revealed disorganized tumoral cytoarchitecture and confirmed the presence of cells representing all three germ layers, such as ectodermal secretory epithelium, mesodermal cartilage and endodermal gut epithelium derivatives (**Fig. 2E**). Together these findings confirmed the *in vitro* and *in vivo* pluripotency of the iPSCs derived from adult NSCs.

### Expression of imprinted genes is altered during the reprogramming of NSCs into iPSCs

To investigate the transcriptional changes associated to the reprogramming process, the transcriptome of both adult NSCs and derived iPSCs was profiled with RNA-seq (GSE282749). The principal component analysis (PCA) clearly distinguished iPSCs from NSCs (**Fig. 3A**). Comparison of NSCs versus iPSCs unveiled 11,986 differentially expressed genes, representing more than 37% of all genes expressed in NSCs (**Fig. 3B,C** and **S3A**). Among these alterations, 5,633 genes were downregulated (17,4% of all genes expressed in NSCs) while 6,353 were upregulated (19,6%) in iPSCs compared to adult NSCs (**Fig. 3C** and **S3A**). Gene Ontology (GO) terms analysis showed several biological processes altered in iPSCs compared to NSCs (**Fig. S3B**). Among upregulated genes, processes such as nucleotide metabolism, mitochondrial gene expression and translation or biogenesis of ribonucleoprotein complexes were enriched; while downregulated genes showed enrichment in categories such as forebrain development, neurogenesis and signal transduction (mainly Wnt signalling) (**Fig. S3B**). As expected, RNA-seq analysis confirmed the upregulation of key pluripotency-associated genes in iPSCs compared to NSCs, including *Oct4, Zfp42,* and *Nanog* (**Fig. 3D**). Conversely, neural-specific genes such as *Nestin, Zic1, and Olig2* were downregulated (**Fig. 3D**).

**FIGURE 3.**
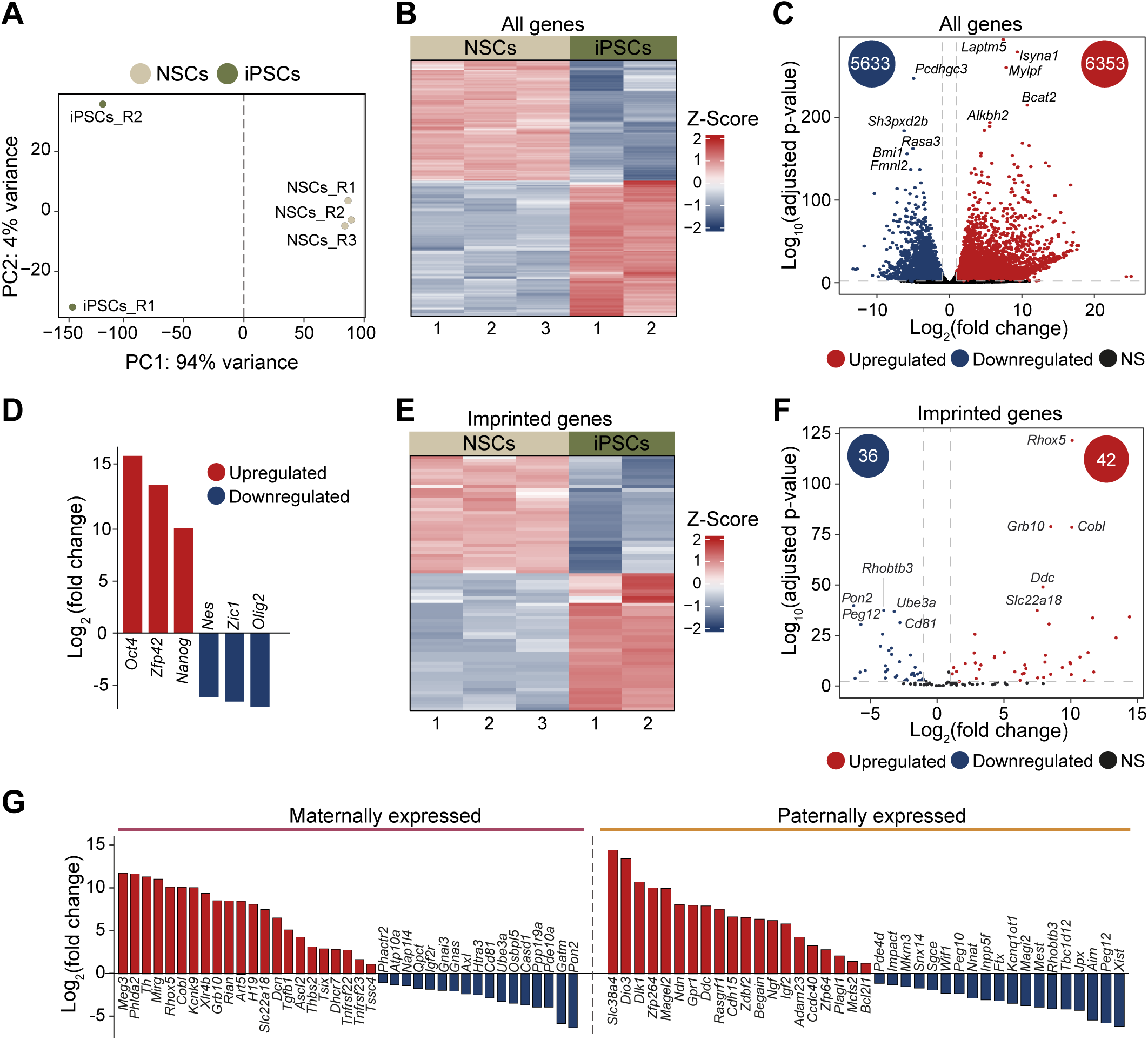
Expression of imprinted genes in adult NSCs is regulated during the reprogramming process. **(A)** Principal component analysis (PCA) generated with top 500 most variable genes obtained from RNA-seq of 3 NSCs and 2 iPSCs cultures, using normalized (variance stabilized transformation) gene counts. It shows principal components 1 and 2. **(B)** Heatmap showing the scaled expression (Z-score) of all differentially expressed genes (DEGs) obtained comparing gene expression levels of iPSC to adult NSCs cultures. **(C)** Volcano plot for all expressed genes based on RNA-seq data. The number of significantly downregulated (blue) and upregulated (red) genes in iPSCs compared to NSCs is indicated inside the circles. **(D)** RNA-seq log_2_(fold change) of gene expression in iPSCs compared to NSCs for the three pluripotency-related genes *Oct4, Zfp42* and *Nanog* and for the three neural genes *Nestin (Nes)*, *Zic1* and *Olig2*. **(E)** Heatmap showing the scaled expression (Z-score) of the imprinted DEGs in iPSCs compared to NSCs. **(F)** Volcano plot for all expressed imprinted genes based on RNA-seq data. The number of significantly downregulated (blue) and upregulated (red) genes in iPSCs compared to NSCs is indicated inside the circles. **(G)** Representation of the log_2_(fold change) of all imprinted DEGs in iPSCs compared to NSCs. Maternally (left panel) and paternally (right panel) expressed genes are shown separately.

Some alterations of genomic imprinting have been observed during reprogramming of somatic cells (Perrera & Martello, 2019; Takikawa *et al*, 2013; Arez *et al*, 2022). These changes have an impact on stem cell plasticity suggesting that genomic imprinting may be a mechanism employed to modulate gene dosage to control stem cell potential (Ferrón *et al*, 2011; Perez *et al*, 2016). Therefore, we next focused on the study of the regulation of imprinted genes during the reprogramming process using the RNA-seq data obtained in adult NSCs and iPSCs. We identified 78 imprinted genes that were differentially expressed between iPSCs and NSCs, representing 60% of all analysed imprinted genes (**Fig. 3E,F** and **S3A**). Among them, similar changes in both paternally and maternally expressed genes were observed (**Fig. 3G**). Imprinted genes expression changes were validated by qPCR in NSCs and iPSCs (**Fig. S3C**), confirming that the acquisition of a pluripotent state also associates with significant transcriptional changes of imprinted genes.

### Acquisition of pluripotency in adult NSCs requires global DNA hypomethylation

DNA methylation represents one of several epigenetic mechanisms employed by cells to regulate gene expression during cell fate decisions (Parry *et al*, 2021). Moreover, previous studies have shown that changes in DNA methylation patterns are essential for successful cell reprogramming (Hochedlinger & Jaenisch, 2015; Lee *et al*, 2014; Takahashi & Yamanaka, 2006). In order to characterize methylation changes at a global level, immunoprecipitation of methylated DNA using an antibody against 5-methylcytosine (5mC) followed by high-throughput sequencing (MeDIP-seq) was performed in adult NSCs and NSC-derived iPSCs (GSE282748) (**Fig. S4A**). A PCA of the results showed clear segregation between NSCs and iPSCs (**Fig. 4A**). Previous studies have reported that iPSCs exhibit lower levels of DNA methylation than somatic cells, highlighting DNA demethylation as a crucial chromatin feature for achieving pluripotency (Lee *et al*, 2014). Consistent with this, the MeDIP-seq analysis revealed genome-wide hypomethylation during the acquisition of the pluripotent state (**Fig. 4B,C** and **S4B**). Among the regions with altered methylation status, 97% displayed hypomethylation (11,865 regions), while only 3% were hypermethylated (362 regions) (**Fig. 4B**). This global hypomethylation was particularly pronounced at promoters and transcription start sites (TSS), as demonstrated by overlapping methylation changes with the 15 chromatin states model (Vu & Ernst, 2023) (**Fig. S4C**). To explore the relationship between transcriptional changes and DNA methylation, RNA-seq and MeDIP-seq data were integrated (**Fig. 4C**). The average methylation signal at the TSS revealed that upregulated genes exhibited a sharper methylation drop in iPSCs compared to the downregulated ones (**Fig. 4D**). Furthermore, the overlap of DMRs with promoter regions showed significant greater hypomethylation in upregulated genes compared to downregulated and unaltered genes (**Fig. S4D**). Gene ontology (GO) analysis of the intersected data revealed that upregulated genes with hypomethylated promoters were enriched in biological processes related to cell division and metabolism, while downregulated genes with hypermethylated promoters were associated with neural development (**Fig. 4E**).

**FIGURE 4.**
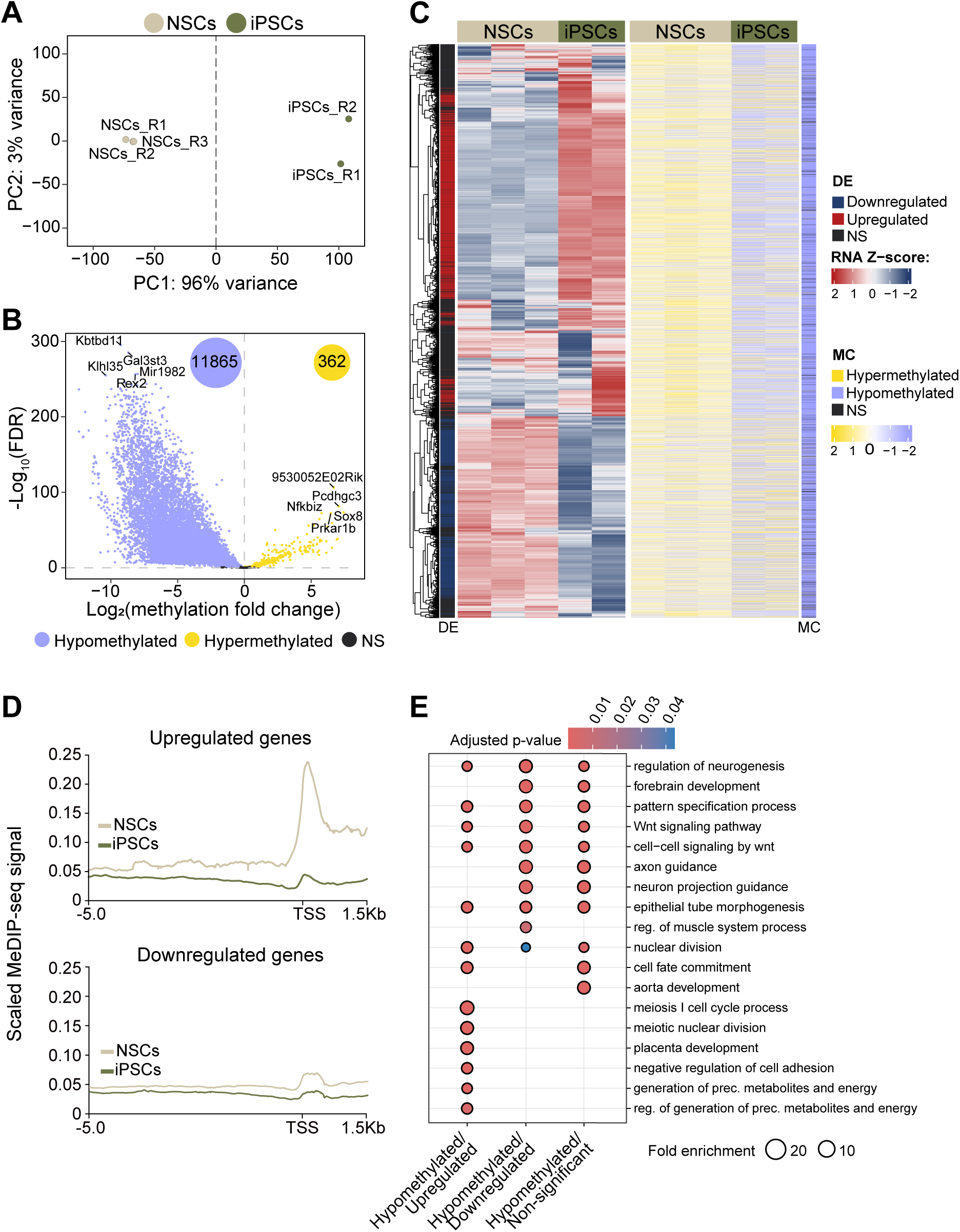
A global hypomethylation is observed in iPSCs genome compared to adult NSCs. **(A)** PCA from MeDIP-seq of 3 NSCs and 2 iPSCs cultures based on normalized (variance stabilized transformation) counts. **(B)** Volcano plot showing the differential methylation signal between iPSCs and NSCs. The number of significantly hypomethylated (purple) and hypermethylated (yellow) regions in iPSCs compared to NSCs is also displayed. **(C)** Heatmap representing both normalized expression and methylation of the whole genome in NSCs and iPSCs. Differential expression (DE) is shown in red (upregulation) and blue (downregulation). Methylation changes (MC) are shown in yellow (hypermethylation) and purple (hypomethylation). Non-significant (NS) changes are shown in black. **(D)** Distribution of 5mC methylation signal around transcription start sites (TSS) of upregulated genes (upper panel) and dowregulated genes (lower panel). **(E)** Gene ontology (GO) analysis of biological processes for gene groups identified with RNA-seq and MeDIP-seq data intersection. Dot size denotes the fold enrichment of each ontology over the background. DMR: differentially methylated region.

### IG-DMR escapes global DNA hypomethylation during acquisition of pluripotency in adult NSCs

Given the variability in the extent and nature of the methylation changes at ICRs reported during pluripotency, we analysed the methylation dynamics across all described imprinted clusters (Santini *et al*, 2021) during adult NSCs reprogramming. Consistent with the genome-wide trends we previously observed, hypomethylation occurred in both germline and somatic ICRs in iPSCs compared to adult NSCs (**Fig. 5A**). Specifically, MeDIP-seq analysis revealed that 24 out of the 25 germline ICRs (96% of all described gDMRs) and 7 out of the 13 somatic ICRs (54% of all described sDMRs) were hypomethylated in iPSCs relative to NSCs (**Fig. 5A; Tables S1** and **S2**). Notably, the IG-DMR previously reported to be hypermethylated in adult NSCs (Ferrón *et al*, 2011), retained its hypermethylated status in iPSCs (**Fig. 5A; Tables S1** and **S2**). To validate these findings, bisulfite DNA treatment followed by pyrosequencing was performed, confirming the hypomethylation of most ICRs and the associated loss of imprinting in these clusters (**Fig. 5B**). Furthermore, the hypermethylation of the IG-DMR in both NSCs and iPSCs was confirmed (**Fig. 5B**), demonstrating that this ICR resists the wave of hypomethylation occurring during iPSC reprogramming.

**FIGURE 5.**
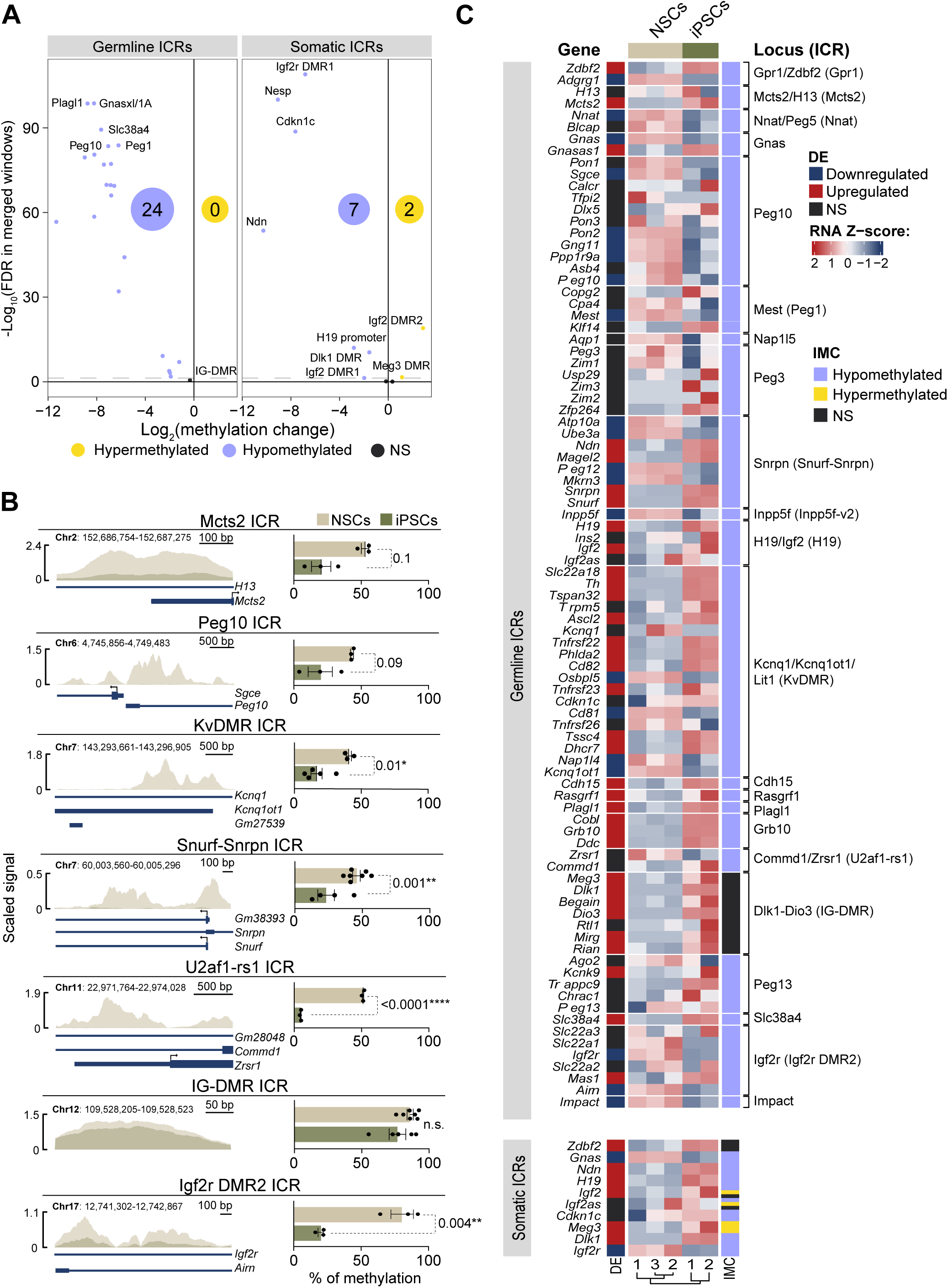
Alterations of methylation and gene expression are observed in different imprinting clusters. **(A)** Volcano plot representing the methylation change between iPSCs and NSCs of different imprinting control regions (ICRs). The number of hypomethylated (purple) and hypermethylated (yellow) DMRs is shown. **(B)** MeDIP-seq methylation signal representation at ICR loci in NSCs and iPSCs (left panels). DNA methylation quantification by pyrosequencing at specific ICRs in NSCs and iPSCs are shown (right panels). Genomic positionss based on the Genome Reference Consortium Mouse Build 38 (GRCm38/mm10) are also indicated. **(C)** Heatmap representing both normalized expression (DE) and methylation (IMC) changes of imprinted clusters. Imprinted genes are cluster-organized, showing the methylation status of each ICR in iPSCs compared to NSCs. In bar plots, error bars show s.e.m.

To investigate whether the observed changes in DNA methylation at ICRs were associated with alterations in imprinted genes expression, we compared transcriptomic data with the MeDIP-seq results in iPSCs and NSCs (**Fig. 5C**). We found that 25% of the changes in imprinted gene expression correlated with the loss of methylation at ICRs in iPSCs (**Fig. 5C**). Focusing on the IG-DMR, which regulates several genes within the *Dlk1-Dio3* cluster (Ferrón *et al*, 2011; Montalbán-Loro *et al*, 2021) (**Fig. S5A**), we observed that the expression levels of *Rtl1* remained unchanged in iPSCs compared to adult NSCs, as expected (**Fig. 5C**). Surprisingly, other genes within the cluster, such as *Begain*, *Dlk1, Meg3,* and *Dio3* were upregulated in iPSCs (**Fig. 5C**). The upregulation of *Dlk1* and *Meg3* could not be attributed to methylation changes at their somatic DMRs, as *Dlk1DMR was* hypomethylated and *Meg3DMR* remained hypermethylated in iPSCs (**Fig. 5C**). Interestingly, while the promoter region of some genes in the cluster (*Meg3* and *Rtl1*) showed no detectable methylation signal (**Fig. S5B**), significant hypomethylation was observed at the promoters of the upregulated genes *Begain*, *Dlk1,* and *Dio3* (**Fig. S5B**). This suggests an additional transcriptional mechanism regulating gene dosage within this cluster. Together, these findings demonstrate that although the IG-DMR resists the widespread hypomethylation associated with reprogramming, the altered expression of genes within the *Dlk1-Dio3* locus arises from transcriptional mechanisms independent of genomic imprinting.

### TET3-mediated transcriptional regulation of *Trim28* preserves IG-DMR hypermethylation in iPSCs

We have demonstrated that the IG-DMR resists the wave of hypomethylation occurring in most ICRs in during the acquisition of a pluripotent state. TET enzymes are dioxygenases that convert 5mC to 5hmC resulting in the removal of methylation marks (Wu & Zhang, 2017) and their role has been demonstrated to be critical for iPSCs reprogramming (Sardina *et al*, 2018). qPCR analysis showed that *Tet3* is the most abundant member of the *Tet* dioxygenases in NSCs isolated from the adult SVZ (Montalbán-Loro *et al*, 2019) (**Fig. 6A** and **S6A**). To investigate the potential role of TET3 in the demethylation process of ICRs during the acquisition of a pluripotent state, a murine genetic model with a conditional deletion of Tet3 in Gfap-expressing cells (mainly NSCs and astrocytes) was generated (see methods). *Tet3* expression ablation was confirmed in NSCs isolated from the SVZ of *Tet3-deficient* (*Gfap-Tet3^KO^*) compared to wild-type (*Gfap-Tet3^WT^*) (**Fig. S6B**). To induce reprogramming, *Gfap-Tet3^KO^* and *Gfap-Tet3^WT^* derived NSCs were co-transduced with the combination of *Oct4, Klf4* and mCherry-encoding retroviral supernatants as previously described (**Fig. 1B** and **S6C-E**). To confirm the full reprogramming of *Tet3KO* NSCs into iPSCs, pluripotency markers and silencing of neural genes were analysed by qPCR in several iPSCs clones (**Fig. 6B** and **S6F**). Although the expression levels of the pluripotency-related gene *Zfp42* and the neural gene *Olig2* were indistinguishable between wild-type and *Tet3*-deficient iPSCs (**Fig. S6F**), an incomplete upregulation of *Nanog* and downregulation of the neural marker *Nestin* were observed in *Tet3*-deficient iPSCs compared to wild-type (**Fig. 6B**). These results suggest that absence of TET3 impairs acquisition of full pluripotency of adult NSCs.

**FIGURE 6.**
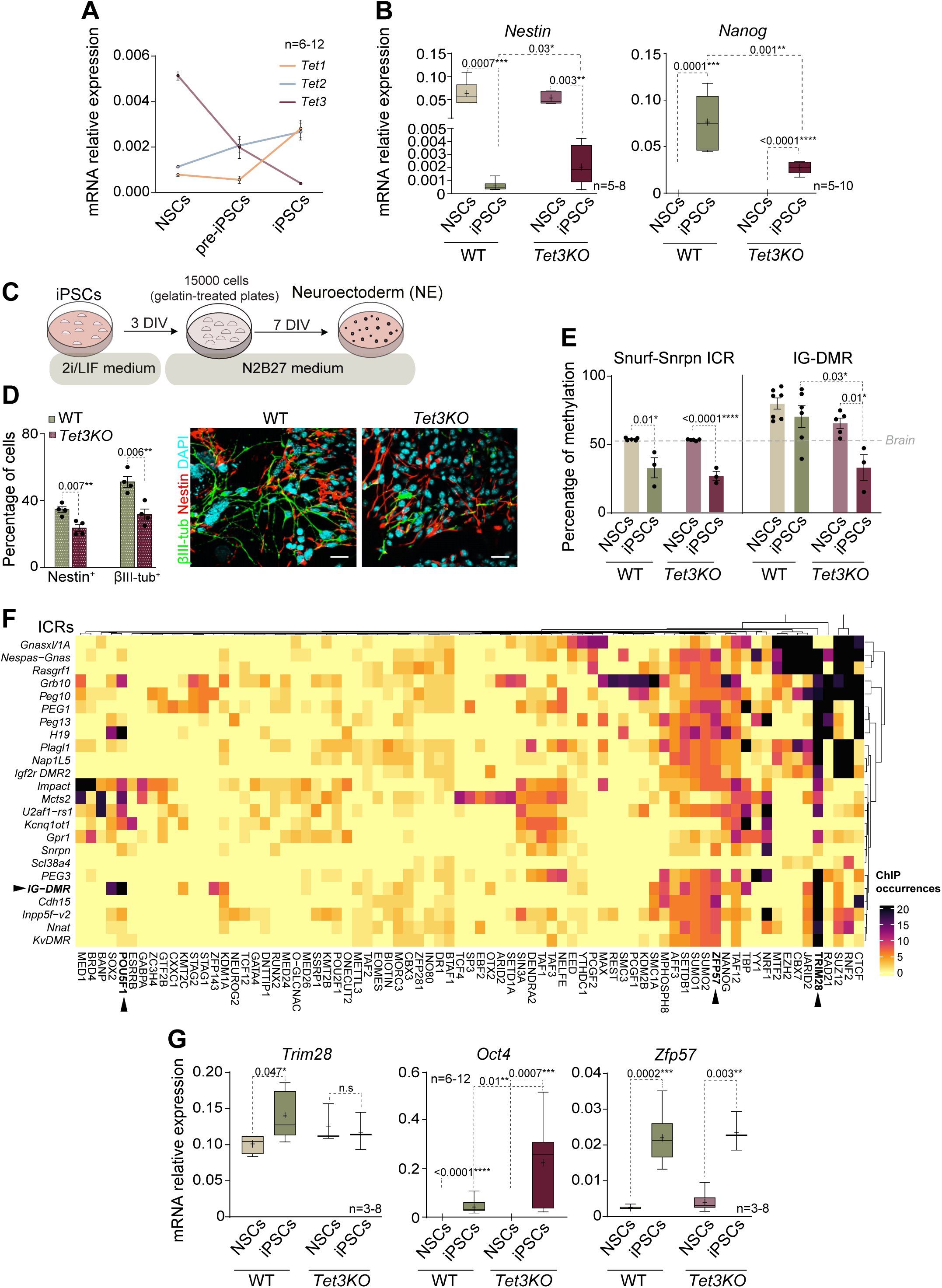
TET3 mediates IG-DMR methylation protection by regulating *Trim28* and *Oct4* gene expression. **(A)** qPCR quantification of *Tet1*, *Tet2* and *Tet3* in NSCs, pre-iPSCs and iPSCs. **(B)** qPCR quantification of the neural marker *Nestin* and the pluripotency marker *Nanog* in wild-type (WT) and *Tet3*-deficient (Tet3KO) NSCs and iPSCs. **(C)** Schematic of the protocol used to differentiate iPSCs into neuroectoderm (NE). iPSCs are disaggregated and re-plated in gelatin-treated plates at a 1.5 ×10^4^ cells/cm^2^ density in N2B27 supplemented medium. Seven days after cells are analyzed. **(D)** Percentage of Nestin and βIII-tubulin positive cells in iPSCs and NE cultures of both genotypes (left panel). Immunocytochemistry images of Nestin (red) and βIII-tubulin (green) in NE cultures of both genotypes (right panel). **(E)** Quantification by pyrosequencing of the percentage of methylation Snurf-Snrpn DMR and IG-DMR in NSCs and iPSCs from both WT and Tet3KO cultures. Gray dashed line indicates the percentage of methylation in control brain samples. **(F)** Heatmap showing ChIP-seq peaks of transcription factors and other proteins in pluripotent stem cells (ChIP-Atlas). Colours correspond to the number of occurrences (number of peaks) each protein binds to different ICRs. **(G)** qPCR analysis of *Trim28*, *Oct4* and *Zfp57* in NSCs and iPSCs from WT and Tet3KO mice. *Gapdh* was used as a housekeeping gene for qPCR analysis. DAPI was used to counterstain DNA. P-values and number of samples are indicated. In box and whiskers plots, the mean is indicated as + and whiskers represent the maximum and minimum values. In bar plots, error bars show s.e.m.

To determine whether *Tet3* deficiency affects the pluripotency of iPSCs, we induced EB formation using the hanging drop method (**Fig. 2A**). Expression analysis of differentiation genes in *Tet3KO* EBs revealed a significant upregulation of mesoderm (*Kdr1)*, endoderm (*Foxa)* and ectoderm (*Cer1)* markers (**Fig. S6G**). Consistently, immunocytochemistry for Brachyury (mesoderm), α-fetoprotein (endoderm) and βIII-tubulin (neuroectoderm) confirmed that *Tet3KO* iPSCs could give rise to cells from all three germ layers (**Fig. S6H**). To further evaluate the differentiation potential of *Tet3-* deficient iPSCs, we directed their differentiation into neuroectoderm (NE) using LIF-free medium with N2 and B27 serum-free supplements (**Fig. 6C**). qPCR analysis of neural markers *Nestin* and *Tubb3* showed that *Tet3KO* iPSCs failed to achieve physiological expression levels of these genes (**Fig. S7A**). Additionally, immunocytochemistry of *Tet3KO* NE cultures showed a significant reduction in the percentage of Nestin^+^ and βIII-tubulin*^+^* cells compared to wild-type iPSCs (**Fig. 6D**). These findings demonstrate that TET3 plays a critical role in the acquisition of naive pluripotency during the reprogramming of adult NSCs and is essential for proper neural differentiation.

To investigate whether the loss of differentiation capacity in *Tet3-deficient* iPSCs correlates with altered methylation levels at ICRs, we performed bisulfite conversion followed by pyrosequencing of the Snrpn-DMR in wild-type and *Tet3KO* iPSCs. NSCs from both genotypes were also analysed. As previously shown, wild-type iPSCs exhibited hypomethylation at the Snrpn-DMR compared to NSCs (**Fig. 6E**). Similarly, *Tet3*KO iPSCs also showed hypomethylation of this DMR (**Fig. 6E**), indicating that TET3 does not regulate the methylation status of this specific DMR in iPSCs. Notably, the hypermethylation pattern observed at the IG-DMR in wild-type NSCs and iPSCs confirmed its protection during reprogramming (**Fig. 6E**). However, a significant loss of methylation was observed in *Tet3KO* iPSCs (**Fig. 6E**), demonstrating that TET3 is crucial for maintaining IG-DMR methylation levels throughout the reprogramming process.

To elucidate the specific role of TET3 in safeguarding IG-DMR methylation during reprogramming, we performed an *in silico* analysis of TET3 binding using chromatin immunoprecipitation followed by deep sequencing (ChIP-seq) data from the ChIP-Atlas database (Zou *et al*, 2024). The analysis showed that TET3 does not directly bind to the IG-DMR (**Fig. S7B**). This finding led us to hypothesize that TET3 might protect IG-DMR methylation in iPSCs by regulating the expression of genes encoding proteins that bind this ICR. To test this hypothesis, we used the ChIP-Atlas database to identify proteins binding to the IG-DMR (Zou *et al*, 2024). Among these, TRIM28 (also known as KAP1 or TIF1-beta) was identified as a protein with high binding occurrences at the IG-DMR (**Fig. 6F**). TRIM28 interacts with the KRAB domain zinc finger protein ZFP57, which serves as a scaffold for recruiting multiple epigenetic factors (Messerschmidt, 2012). Consistent with this, our analysis also identified ZFP57 as an IG-DMR-binding protein (**Fig. 6F**). Additionally, OCT4, which has been implicated in driving ICR hypomethylation in post-implantation embryos (Zimmerman *et al*, 2013), was also found to bind the IG-DMR (**Fig. 6F**). qPCR analysis revealed upregulation of *Trim28*, *Oct4* and *Zfp57* in wild-type iPSCs compared to NSCs (**Fig. 6G** and **S7C**). Notably, *Trim28* upregulation was not observed in *Tet3KO* iPSCs (**Fig. 6G**), while *Oct4* levels were significantly higher in *Tet3KO* iPSCs than in wild-type iPSCs (**Fig. 6G**) and remained elevated even after differentiation into neuroectoderm (**Fig. S7D**). In contrast, *Zfp57* expression was unaffected in *Tet3KO* iPSCs (**Fig. 6G**). These observations align with ChIP-seq data showing TET3 binding at the promoters of *Trim28* and *Oct4* (**Fig. S7E**), suggesting that TET3 regulates these genes. Together, these findings reveal a non-canonical role for TET3 in protecting the IG-DMR from global hypomethylation during reprogramming potentially modulating *Trim28* and *Oct4* expression (**Fig. S7F**).

## Discussion

Our study uncovers a novel mechanism in which the dioxygenase TET3 plays a pivotal role in maintaining methylation levels during the reprogramming of adult NSCs to iPSCs. Reprogramming with the transcription factors *Oct4* and *Klf4* induces global gene expression changes and a widespread loss of DNA methylation, including at ICRs. Remarkably, we found that the IG-DMR, the ICR regulating the imprinted *Dlk1-Dio3* cluster on mouse chromosome 12, is uniquely protected from this demethylation wave. This protection depends critically on TET3, which acts indirectly through transcriptional regulation of *Trim28* and *Oct4*. Our model suggests that TRIM28 protein binds to the IG-DMR and recruits DNMT enzymes to preserve DNA methylation during reprogramming. However, in the absence of TET3, TRIM28 is not expressed at sufficient levels to protect the IG-DMR, allowing OCT4 to bind and promote its demethylation. Importantly, the loss of TET3 not only disrupts the maintenance of methylation at the IG-DMR but also impairs the differentiation potential of the resulting iPSCs. This suggests that the epigenetic instability caused by TET3 deficiency compromises the ability of iPSCs to properly execute lineage-specific differentiation programs. These findings highlight a previously unrecognized role of TET3 in preserving the epigenetic status of the *Dlk1-Dio3* cluster and ensuring the functional integrity of pluripotent stem cells, emphasizing its importance for successful reprogramming and differentiation.

Through genome-wide expression analysis using next-generation sequencing, we demonstrate that various markers are activated or repressed following reprogramming induction, both in iPSCs and in the adult NSCs of origin. Specifically, as expected, the repression of neural markers such as *Nestin* and *Olig2* correlates with the activation of pluripotency genes like *Oct4*, *Nanog*, and *Zfp42* in iPSCs. Furthermore, a GO analysis reveals that multiple gene sets, associated with key biological pathways, are significantly upregulated or downregulated in iPSCs compared to adult NSCs, confirming that the transition to pluripotency involves broad transcriptomic changes. Our study also highlights that epigenetic remodelling during the reprogramming of NSCs into iPSCs is critical for pluripotency induction. A global change in the DNA methylation is necessary for achieving true pluripotency. Detailed analysis of the methylation profiles of pluripotency-associated genes showed DNA hypomethylation, particularly at the promoter regions, while neural-associated genes exhibited increased methylation, consistent with their repression. Imprinted genes, which are expressed monoallelically from either the maternal or the paternal chromosome (Tucci *et al*, 2019), represent a unique epigenetic subset that poses additional challenges during reprogramming. Consistently, our RNA-seq data show that approximately 60% of all imprinted genes analyzed undergo transcriptional changes in iPSCs derived from NSCs, suggesting that the regulation of imprinted gene expression plays a critical role in both NSC behaviour and the reprogramming process.

DNA methylation at ICRs is essential for both the establishment and maintenance of genomic imprinting during development (Bartolomei & Ferguson-Smith, 2011). Germline DMRs are crucial for initiating monoallelic expression during development, while somatic DMRs play pivotal roles in maintaining imprinting in somatic cells (Ferguson-Smith, 2011). Naïve iPSCs exhibit several features similar to early mammalian embryos, including global genome hypomethylation, which is accompanied by a widespread loss of DNA methylation at imprinted loci (Perrera & Martello, 2019). In line with this, our findings reveal that the majority of ICRs in imprinted clusters are also hypomethylated in iPSCs. These epigenetic alterations correlate with changes in the expression levels of several imprinted genes within these clusters, as observed by RNA-seq, suggesting that the reprogramming process contributes to the regulation of genomic imprinting during the acquisition of pluripotency.

Previous studies from our group have revealed that differential DNA methylation at the IG-DMR on chromosome 12 is not maintained in adult NSCs and niche astrocytes in the adult mouse brain (Ferrón *et al*, 2011). In these cells, the IG-DMR becomes hypermethylated, leading to bi-allelic expression of the *Dlk1* gene, in contrast to the mono-allelic expression observed in embryos (Ferrón *et al*, 2011; Montalbán-Loro *et al*, 2021). Studies from other groups have proposed the *Dlk1-Dio3* cluster as a potential exclusive marker for iPSCs, noting that iPSCs with normal *Dlk1-Dio3* expression generate high-grade chimeras, while those with silenced expression contribute poorly to chimeras (Benetatos *et al*, 2014). Contradictory findings suggest that this locus may not be critical for reprogramming, as they report no significant role for *Dlk1-Dio3* in iPSC formation (Stadtfeld *et al*, 2010; Li *et al*, 2011; Pham *et al*, 2022). We reveal here that the IG-DMR resists the wave of hypomethylation occurring during adult NSC reprogramming, thereby maintaining the original hypermethylated status at this locus in iPSCs. We postulate that gain of methylation in NSCs and further maintenance of IG-DMR hypermethylation in iPSCs may be essential for achieving full pluripotency during reprogramming. We also propose that the methylation level of the *Dlk1-Dio3* cluster may be a more reliable pluripotency marker, suggesting a clear influence of the *Dlk1-Dio3* locus in iPSCs pluripotency.

Many questions remain regarding how mechanistically genomic imprinting is established, maintained and then erased during reprogramming. It is well known that DNA demethylation proceeds through one of two distinct mechanisms: passive loss of 5mC during DNA replication via suppression of DNMT activity or active demethylation by TET (Hill *et al*, 2014; Tahiliani *et al*, 2009). Active demethylation is initiated through the progressive oxidation of 5mC to 5hmC (Ito *et al*, 2011), after which demethylation is achieved (Hashimoto *et al*, 2012). While *Tet1* and *Tet2* expression is induced during iPSC generation, a higher level of expression of *Tet3* in adult NSCs is observed (Montalbán-Loro *et al*, 2019). Here we address the potential role of TET3 in preserving IG-DMR methylation during iPSCs reprogramming by using a murine model with conditional *Tet3* deletion in *Gfap*-expressing cells. Our results confirm that *Tet3*-deficient NSCs showed incomplete reprogramming into iPSCs, with reduced upregulation of *Nanog* and diminished downregulation of *Nestin*, indicating defective pluripotency acquisition. *Tet3KO* iPSCs maintain their ability to differentiate into all three germ layers, however, when directed towards neuroectoderm differentiation, they also fail to achieve proper neural differentiation, suggesting a critical role for TET3 for the acquisition of naïve pluripotency during reprogramming.

Further methylation analysis using pyrosequencing of the Snrpn-DMR revealed no significant differences in methylation between wild-type and *Tet3KO* iPSCs. However, *Tet3KO* iPSCs exhibited a marked loss of methylation at the IG-DMR, indicating that TET3 is necessary to maintain methylation of this ICR during reprogramming. *In silico* analysis of TET3 binding revealed that TET3 does not directly bind to the IG-DMR, but it likely regulates genes encoding proteins that bind this region. In line with this, we identify TRIM28 and ZFP57 as key proteins with significant binding activity at the IG-DMR in iPSCs. These two proteins have been reported to interact to protect methylation imprints at the IG-DMR from both active and passive demethylation during preimplantation development (Alexander *et al*, 2015). Gene expression analysis confirmed that genes encoding for these two proteins are upregulated in iPSCs, whereas *Trim28* expression was not induced in*Tet3KO* iPSCs indicating a direct regulation of this gene by TET3. Finally, we demonstrate that OCT4, a key pluripotency factor known to promote hypomethylation of ICRs in post-implantation embryos (Zimmerman *et al*, 2013), also binds the IG-DMR. *Oct4* levels were upregulated during reprogramming and were significantly higher in *Tet3*-deficient compared to wild-type iPSCs, suggesting that TET3 also regulates the *Oct4* expression. ChIP-seq data showing TET3 binding at the promoters of *Trim28* and *Oct4* indicate that TET3 may transcriptionally regulate these genes, which, in turn, play opposing roles in regulating IG-DMR methylation.

In conclusion, our study reveals that loss of imprinting is observed in iPSCs generated from adult NSCs. This is often regarded as a dysfunctional state that conflicts with the establishment of pluripotency. However, the hypermethylation observed at the IG-DMR in iPSCs already exists in NSCs and is essential for their proper function (Ferrón *et al*, 2011; Montalbán-Loro *et al*, 2021). Therefore, changes in the methylation at some ICRs could be a pivotal feature in transitioning to pluripotency, indicating that its presence is not inherently detrimental but may reflect a necessary step in the epigenetic reorganization of reprogramming. Moreover, our findings uncover a non-canonical role for TET3 in modulating key pluripotency genes like *Oct4* and *Trim28*. We propose that the recruitment of TRIM28 to the IG-DMR contributes to the maintenance of IG-DMR methylation in iPSCs. In this context, where the balance between methylation and demethylation is pivotal, the absence of TET3 may facilitate the aberrant binding of OCT4 to the IG-DMR. This would promote the demethylation of this region, disrupting its protective methylation marks and potentially impairing the proper establishment of pluripotency. Therefore, the role of TET3 in protecting IG-DMR methylation by repressing *Oct4* and thus restricting OCT4 access to this ICR represents a critical aspect of the epigenetic regulation required for successful reprogramming.

## Methods

### Animals and *in vivo* manipulations

The experiments were conducted in C57BL/6 wild-type mice. For the teratoma formation assays, homozygous immunosuppressed *Nude* (NU/J) mice obtained from the Jackson Laboratory were used. For *Tet3* delection in GFAP^+^ NSCs, heterozygous GFAP-cre transgenic animals (6.Cg-Tg(Gfap-cre)73.12Mvs/J) from the Jackson Laboratory were bred to *Tet3^loxp/loxp^* (Montalbán-Loro *et al*, 2019). Expression of Cre-recombinase under the *Gfap* promoter results in a deletion of exon 5 of *Tet3* gene, causing a frame-shift from exon 6 and a premature stop codon in exon 7 of the gene. Animals were genotyped by PCR analysis of DNA as described (Montalbán-Loro *et al*, 2019) and littermates lacking *GFAP-Cre* were used as control mice. All mice were maintained on a C57BL6 background and in a 12 h light/dark cycle with free access to food and water *ad libitum* and according to the Animal Care and Ethics committee of the University of Valencia.

### Neurosphere cultures

Adult 2- to 4-month-old mice were euthanized by cervical dislocation. To initiate each independent culture, the brains of two different animals were dissected and the regions containing the SVZ were isolated from each hemisphere and washed in Earle’s balanced salt solution (EBSS; Gibco). Tissues were transferred to EBSS containing 12 U mL^−1^ papain (Worthington DBA), 0.2 mg mL^−1^ L-cystein (Sigma), 0.2 mg mL^−1^ EDTA (Sigma) and incubated for 20 min at 37°C. Tissue was then rinsed in EBSS, transferred to Dulbecco’s modified Eagle’s medium (DMEM)/F12 medium (1:1 v/v; Life Technologies) and carefully triturated with a fire-polished Pasteur pipette to a single cell suspension. Isolated cells were collected by centrifugation, resuspended in DMEM/F12 medium containing 2 mM L-glutamine (Gibco), 0.6% glucose (Panreac), 9.6 g mL^−1^ putrescine (Sigma), 6.3 ng mL^−1^ progesterone (Sigma), 5.2 ng mL^−1^ sodium selenite (Sigma), 0.025 mg mL^−1^ insulin (Sigma), 0.1 mg mL^−1^ transferrin (Sigma), 2 μg mL^−1^ heparin (sodium salt, grade II; Sigma) and supplemented with 20 ng mL^−1^ epidermal growth factor (EGF; Invitrogen) and 10 ng mL^−1^ fibroblast growth factor (Belenguer *et al*, 2016) (FGF; Sigma). Neurospheres were allowed to develop for 6 days in a 95% air-5% CO_2_ humidified atmosphere at 37 °C. For culture expansion, cells were plated at a relatively high density (75 cell/μl) and maintained for several passages.

### Reprogramming of adult NSCs into iPSCs

To generate iPSCs from adult NSCs, exogenous *Oct4* together with *Klf4* (2F) were used for reprogramming as previously described (Kim *et al*, 2008b). To produce retroviruses expressing *Oct4* and *Klf4*, Platinum-E (Plat-E) retroviral packing cells (Cell Biolabs) were transfected with a plasmid solution containing 1 mL of Opti-MEM^TM^ (Gibco), 60 μL of 1mg mL^−1^ polyethylenimine (PEI, Polysciences) and 20 µg of the retroviral vectors pMXs-*Oct4* (#13366, Addgene), pMXs-*Klf4* (#13370, Addgene) and pMXs-*mCherry* (pMX-2A-CH, designed and kindly provided by Dr. Jose Manuel Torres). After 24 hours, Plat-E culture medium (high glucose DMEM containing 10% foetal bovine serum FBS, 2 mM L-glutamine, 1 μg mL^−1^ Puromycin and 10 μg mL^−1^ Blasticidin) was replaced by NSCs complete medium. Transfection efficiency was checked by mCherry fluorescence in Plat-E cells (**Fig. S1A**). The day after, retrovirus-containing supernatants were collected and filtered with a 0.45 μm nitrocellulose filter. Neurospheres from female mice were grown for two days were transduced with a mixture of these supernatant (SN) as follows (volume *per* plate): 3 mL of *Oct4* SN, 3 mL of *Klf4* SN, 1 mL of *mCherry* SN and 3 ml of fresh NSCs complete medium. A control of infection was made with a mixture containing 7 ml of *mCherry* retrovirus containing medium and 3 mL of fresh complete medium. In order to enhance the efficiency of retroviral infection, retrovirus mixture was supplemented with 4 μg mL^−1^ of polybrene (Sigma). NSCs were then incubated for 14-18 hours at 37°C in a humidified incubator. Infected NSC medium was then replaced with fresh complete medium and neurospheres were allowed to develop for 5 days (**Fig. 1B**). The mouse fibroblast cell line SNL (Cell Biolabs) was used as feeder cells during the reprogramming process. SNL feeder cells were first mitotically inactivated by treatment with 4 μg mL^−1^ of Mitomycin C (Sigma) for 2-4 hours. Plates were treated with 0.1% of gelatin (Sigma) at 37°C for at least 20 min and then mitomyzed SNLs were plated at high density (2.5×10^6^ cell/plate) in gelatine-treated plates (day 7). Five days after transduction, neurospheres were dissociated with Accutase® and 1.5×10^5^ of infected NSCs were re-plated on SNL feeder cells with ESC/LIF medium: Glasgow Minimum Essential Medium (GMEM) containing 15% FBS, 2 mM L-glutamine, 1 mM Sodium pyruvate (Gibco) and 1 μM Leukaemia Inhibitory Factor (LIF). ESC/LIF medium was changed every other day until Stage-Specific Embryonic Antigen-1 (SSEA-1; also known as CD15) positive colonies appeared (pre-iPSCs), checked by staining with StainAlive SSEA-1 Antibody (DyLight 488) (Stemgent®, 1:100 dilution) (**Fig. 1F** and **Fig. S1A**). ESC/LIF medium was replaced with 2i/LIF Neurobasal medium containing B27 supplement; Gibco, 2 mM L-glutamine, 1 mM Sodium pyruvate, 1 mg mL^−1^ Transferrin, 50 μM Insulin, 16 μg mL^−1^ Putrescine, 60 ng mL^−1^ Progesterone, 0.3 μM Sodium selenite, 50 μg mL^−1^ Bovine serum albumin, 1 μM LIF, 1 μM iMEK (Millipore, PD03259Platinum and 3 μM iGSK3 (Millipore, CHIR99021) which is based on dual inhibition (2i) of mitogen-activated protein kinase (MAPK) signalling and glycogen synthase kinase-3 (GSK3) combined with LIF (Silva *et al*, 2008). 2i/LIF medium was changed every two days until well-defined iPSCs colonies appeared (**Fig. 1B**). To establish and expand clonal lines of iPSCs, individual colonies were isolated and plated on gelatine treated plates with 2i/LIF medium. The embryonic stem cell (ESC) line E14Tg2a was used as a pluripotency positive control in the different experiments. ESCs were cultured on gelatine-treated plates and 2 days after plating, cells were treated with Trypsin/EDTA and re-plated following a dilution of 1:5 in ESC/LIF medium.

### Embryoid bodies assays

Embryoid bodies were obtained using the “*hanging drops*” method. iPSCs were treated with Accumax® (Millipore) and resuspended in EB medium: GMEM containing 10% FBS, 2 mM L-glutamine and 1 mM Sodium pyruvate. Several rows of 20 μl drops of a cell suspension at 30 cells/ μL were plated using a multichannel pipet (**Fig. 2A**). Plates were incubated upside-down for 3 days at 37°C in a 5% CO_2_ humidified incubator. Plates were then inverted and EB medium was added. To avoid EB attachment plates were previously treated with 0.4% poly (2-HEMA) solution (Sigma) prepared in Ethanol:Acetone (1:1). EBs were incubated for 4 more days and then plated on gelatine-treated plates for 3 more days before analysis (**Fig. 2A**).

### Neuroectoderm differentiation

Neuroectoderm (NE) differentiation was performed as previously described(Ying *et al*, 2003). Briefly, iPSCs were treated with Accumax® and resuspended in NE culture medium: DMEM/F12 GlutaMAX™ (containing 25 μg/mL insulin, 100 μg mL^−1^transferrin, 6 ng mL^−1^ progesterone, 16 μg mL^−1^ putrescine, 30 nM sodium selenite, and 50 μg mL^−1^ BSA) and Neurobasal (containing 2% B27, 2 mM GlutaMAX™, 0.1% 2-mercaptoethanol, and 1.45% sterile glucose) (**Fig. 6C**). Cells were counted and plated at a density of 1.5 ×10^4^ cells/cm^2^ in gelatine pre-treated plates (**Fig. 6C**). Cultures were maintained in NE culture medium and the medium replenished every other day for a week (**Fig. 6C**).

### Teratoma formation and analysis

To evaluate the capacity of iPSCs to generate teratomas, mouse iPSCs cultures were collected by treatment with Accumax®. iPSCs were washed in PBS and resuspended in PBS supplemented with 30% Matrigel® (Corning®) (Prokhorova *et al*, 2009). Cells were kept on ice and drawn into a 1mL syringe immediately before injection. Approximately 1.5×10^6^ cells resuspended in 200 μL of solution were injected in the dorsolateral area of the subcutaneous space on both sides of the mice back. Teratomas were allowed to develop for 15-20 days when the size of the teratomas was approximately 1.5-2 cm. Mice were sacrificed by cervical dislocation and teratomas were extracted for analysis. For teratoma analysis, samples were fixed in 4% paraformaldehyde (PFA) overnight at 4°C with shaking. Samples were embedded in paraffin and teratoma samples were serially sectioned into 7 μm sections using a microtome (Leica). Slices were stained with haematoxylin and eosin and cell types from the three embryonic layers were identified under the optic microscope (Nikon Eclipse Ni).

### Karyotype of iPSCs

To perform the karyotype analysis, cell division was inhibited using 0.6 μg mL^−1^ of KarioMAX® Colcemid (Gibco) at 37°C. After 2 hours, culture medium was removed and 0.85% sodium citrate, previously warmed at 37 °C, was added. A cell scraper (Biofil®) was used to raise the cells. Cell suspension was transferred to a 15 mL conical tube and incubated at 37°C for 15 min. After that, 10 drops of cold Carnoy fixative (methanol-acetic acid, 3:1) were added to the suspension and softly mixed using a Pasteur pipette. Samples were washed several times with 5 ml of cold Carnoy solution and, after centrifugation (10 min, 300*g*), pellets were resuspended in 2 drops of Carnoy fixative. Cells extensions were made in microscope slides followed by heat fixation. Samples were stained with Leishmańs stain (Sigma). The number of chromosomes was determined under the optic microscope (Nikon Eclipse Ni).

### Immunocytochemistry and alkaline phosphatase (AP) staining

NSCs, iPSCs and EBs were fixed for staining with 4% PFA in 0.1M PBS for 15 min and immunocytochemistry performed as previously described (Belenguer *et al*, 2016). Primary and secondary antibodies and dilutions used are listed in **Table S3** and **Table S4** respectively. DAPI (1 μg mL^−1^) was used to counterstain DNA. Samples were photographed and analysed using an FV10i confocal microscope (Olympus). Alkaline phosphatase detection method was used in reprogrammed cells to check the presence of iPSCs after one month in 2i/LIF medium on *feeders*. Cells were fixed with cold methanol for 2 min and washed three times with 0.1M tris-HCl pH 8.5 buffer. Samples were incubated with the “*staining solution*” which contained 0.1 mg mL^−1^ naphtol phosphate (Sigma), 0.5% Dimethylformamide (Sigma) and 0.6 mg mL^−1^ Fast Red Salt (Sigma) in 0.1M tris-HCl pH 8.5. When red precipitate appeared, cells were washed with 0.1M tris-HCl and distilled water. Finally, the different plates were photographed using a dissection microscope.

### Expression studies

RNAs were extracted with RNAeasy Mini Kit (Qiagen) including DNase treatment, following the manufacturer’s guidelines. For qPCR, 1 μg of total RNA was reverse transcribed using random primers and RevertAid H Minus First Strand cDNA Synthesis kit (Thermo Scientific), following standard procedures. Thermocycling was performed in a final volume of 10 μl, containing 4-10 ng of cDNA sample and the reverse transcribed RNA was amplified by PCR with appropriate Taqman probes (**Table S5**). qPCR was used to measure gene expression levels relative to *Gapdh*, which expression did not differ among the groups. qPCR reactions were performed in a Step One Plus cycler with Taqman Fast Advanced Master Mix (Applied Biosystems). In case of using SYBR green, thermocycling was also performed in a final volume of 10 μL, containing 4-10 ng of cDNA sample, 0.2 μM of each primer (**Table S6**) and SYBR® Premix ExTaq^TM^ (Takara) according to the manufacture instructions, using ROX as a reference dye. A standard curve made up of doubling dilutions of pooled cDNA from the samples being assessed was run on each plate, and quantification was performed relative to the standard curve.

### RNA-seq

RNA was isolated using TRIzol® following manufacturer’s instructions. Briefly, 1 mL of reagent was added per 5-10×10^6^ cells for lysis during 20 min at room temperature (RT). Then, 100 μL of chloroform was added to samples and incubated at RT for 10 min and centrifuged at 12,000*g* centrifugation at 4°C for 10 min. For RNA precipitation, aqueous phase was mixed with 500 μL of isopropanol and incubated for 5 min. Samples were centrifuged 8 min at 12,000*g* at 4 °C. RNA pellet was washed in 1 mL of 75% ethanol, vortexed and centrifuged 5 min at 7500*g* at 4 °C. Then, RNA pellet was resuspended in RNase-free water and stored at −80°C until use. Library preparation and high-throughput sequencing were performed by the Central Service for Experimental Research (SCSIE) at the University of Valencia. Libraries were generated from triplicated biological samples per condition using the Illumina TruSeq stranded mRNA Sample Preparation Kit v2 following the manufacturer’s protocol and sequenced using Illumina NextSeq 500. Read quality was assessed with *FastQC*. Expression was quantified at gene level with *salmon* (Patro *et al*, 2017) in pseudomapping mode, with automatic library detection (-l A) and sequence bias correction (--seqBias) using the Gencode release M23 as reference (Frankish *et al*, 2020). Gene expression quantification was imported into R with package *tximeta* (Love *et al*, 2020) and differential expression analysis was performed with DESeq2 (Love *et al*, 2014). Gene ontology (GO) analysis was conducted with R package clusterProfiler (Wu *et al*, 2021). *Ggplot2* was used for visualizations and *dplyr*, *tibbl*e and *tidyr* packages for data wrangling (Wickham *et al*, 2019). Heatmap visualizations were performed with Complex Heatmap package (Gu *et al*, 2016) and scaled variance stabilized counts.

### MeDIP-seq

DNA was extracted with DNeasy Blood and Tissue Kit (Qiagen) following the manufacturer’s instructions. Samples were eluted in 100 μLof elution buffer and DNA concentration was measured using a Nanodrop 1000. MeDIP-seq protocol was adapted from Taiwo *et al*., 2012 (Taiwo *et al*, 2012). For immunoprecipitation, 3 μg of DNA were sonicated to obtain 150-200 bp fragments. Sonication efficiency was checked by capillary electrophoresis (Bioanalyzer, Agilent). DNA libraries were prepared using NEBNext® Ultra™ II FS DNA Library Prep Kit for Illumina (New England Biolabs). For MeDIP, 1.5 μg of DNA was diluted in TE buffer (10 mM tris-HCl, 1 mM EDTA, pH 7.5) and denatured for 10 min at 99°C. Non-specific interactions were blocked by adding 20 μl of 10x IP buffer (100 mM Na-Phosphate pH 7.0, 0.5% tritonX-100) and 100 μL of 5% skimmed milk buffer in 2 M NaCl. Then 2 μg of anti-5mC antibody (Diagenode, cat. no. C15200006) were added and incubated for 2 h at 4°C with rotation. In parallel, 11 μl per sample of Dynabeads® M-280 sheep anti-mouse IgG (Thermo Fisher, cat. no. 11201D) were blocked with 500 µl PBS-BSA (1 mg mL^−1^ BSA in 0.1 M PBS) for 2 hours at 4°C with rotation. After incubations, beads were collected in a magnetic rack, re-suspended in the original volume (11 μL) with 1x IP buffer (10 mM Na-Phosphate pH 7.0, 0.05% tritonX-100, 1 M NaCl) and added to the DNA samples, which were incubated overnight at 4°C with rotation. The day after, beads were collected using a magnetic rack and the supernatant (unbound fraction) was transferred to a fresh tube. Beads were washed three times with 500 µl of 1x IP buffer for 10 min with rotation at 4°C. After the final wash, bound and unbound fractions were treated with 0.3 mg/mL of Proteinase K (Roche) in digestion buffer (50 mM Tris-HCl pH 8.0, 10 mM EDTA, 0.5% SDS) and incubated at 55 °C for 30 min on a shaking heating block. Samples were purified using MiniElute PCR purification kit (Qiagen) and eluted in 10 μl of elution buffer. To calculate 5mC enrichment in the bound fraction, quantitative PCRs for unmethylated and methylated regions were done from bound and unbound fractions (**Fig. S4A**). Enrichment should be of at least 25x, specificity should be more than 95% and unmethylated recovery should be less than 1% (**Fig. S4A**). Samples were sequenced in a HiSeq2000 (Illumina, Inc) instrument. Primers used to evaluate MeDIP efficiency and specificity are provided in Supplementary Table S7.

MeDIP-seq reads were pre-processed using MEDUSA pipeline (Wilson *et al*, 2012) with default settings and the mm10 UCSC reference genome. Signal tracks were obtained with *Deeptools* suite (Ramírez *et al*, 2014), with the bamCoverage function, applying the following parameters: --scaleFactor X --binSize 1 --blackListFileName -- minMappingQuality 20. The blacklist file was sourced from Boyle Lab Github repository (Amemiya *et al*, 2019). The scaling factor was determined using the normalization factor obtained from counting reads in 10Kb bins, followed by TMM normalization using the *edgeR::calcNormFactors* function (Robinson *et al*, 2009). Biological replicates were combined using the *bigwigAverage* function from Deeptools, averaging the methylation signal. Read counting, CpG density normalization, and differential methylation analysis were performed using the *MEDIPS* R package (Lienhard *et al*, 2014). A sliding window of 100 bp was applied, and a minimum read depth of 10 across all samples was required for inclusion in the analysis. Differential methylation was modeled with the *edgeR* (Robinson *et al*, 2009) implementation in the *MEDIPS* package, using TMM normalization and multiple testing correction via the Benjamini-Hochberg method. Principal component analyisis (PCA) was performed using the *DESeq2* package (Love *et al*, 2014). To ensure robust signal detection, we retained the top 5% percentile windows with the highest signal and discarded those that did not overlap a CpG island or a CpG shore (obtained from UCSC mm10 repository). Adjacent windows were merged by calculating the harmonic mean of p-values using the extraChIPs::mergeByHMP with the following parameters: merge_within = 1, p_adj_method = fdr, alpha = 0.01. Finally, windows were annotated to genes using the *ChIPseeker* package (Yu *et al*, 2015). For ICR analysis, we used previously defined ICRs (Santini *et al*, 2021), completed with additional somatic ICR locations identify in our laboratory (**Tables S1** and **S2**). We extracted windows overlapping ICR genomic regions prior to CpG filtering using the R package *plyranges* and the *join_overlap_inner* function (Lee *et al*, 2019a). Adjacent windows were merged, as described previously, with the merge_within parameter set to 300. Gene ontology (GO) analysis, data wrangling and visualization were performed with the aforementioned packages. Metagenes were generated by combining the *ComputeMatrix* reference-point and *plotProfile* functions from the *Deeptools* suite, which represent the methylation signal at nucleotide resolution (--bin Size 1) within a −5000/1500 bp window (-a 1500, -b 5000) around the TSS (--reference Point TSS). Signal tracks were visualized using SparK (Kurtenbach & Harbour, 2019), with averaged the scaled methylation signal across biological replicates over the selected ICR genomic regions. Chromatin states were obtained from Vu and Ernst 2023 (Vu & Ernst, 2023) and overlapped using *plyranges* and *join_overlap_inner* function (Lee *et al*, 2019b).

### DNA methylation analysis by pyrosequencing

DNA methylation level was quantified using bisulfite conversion and pyrosequencing. The DNA was bisulfite-converted using EZ DNA Methylation-GoldTM kit (Zymo research) in accordance with the manufacturés protocol. Specifically, for the different DMRs, bisulfite-converted DNA was amplified by PCR with specific primer pairs (**Table S8**). PCRs were carried out in 20 μL, with 2U HotStar Taq polymerase (Qiagen), PCR Buffer 10x (Qiagen), 0.2 mM dNTPs and 400 mM primers. PCR conditions were: 96 °C for 5 min, followed by 39 cycles of 94 °C for 30 s, 54 °C for 30 s and 72 °C for 1 min. For pyrosequencing analysis, a biotin-labelled primer was used to purify the final PCR product using sepharose beads. The PCR product was bound to Streptavidin Sepharose High Performance (GE Healthcare), purified, washed with 70% ethanol, denatured with 0.2 N NaOH and washed again with 10 mM tris-acetate. Pyrosequencing primer (400 mM) was then annealed to the purified single-stranded PCR product and pyrosequencing was performed using the PyroMark Q96MD pyrosequencing system using PyroMark® reactives (Qiagen).

### Statistical analysis

All statistical tests were performed using the GraphPad Prism Software, version 7.00 for Windows. Data were first tested for normality using Shapiro-Wilk test. The significance of differences between groups was evaluated using appropriate statistical tests for each comparison. For data that passed normality tests: a paired t-test was used when comparing only two groups (applying Welch’s correction in case the standard deviation of groups is different); and one-way ANOVA followed by Šídák’s post-hoc test was applied for comparing three or more groups. For data groups that did not pass normality: Mann–Whitney nonparametric test was performed when comparing only two groups and Kruskal-Wallis test followed by Dunn’s post-hoc test when more than two groups were analysed. When comparisons were performed with relative values (percentages), data were previously normalized by using arcsin root transformation. Values of P<0.05 were considered statistically significant. Data are presented as the mean ± standard error of the mean (s.e.m.) and the number of experiments performed with independent cultures or animals (*n*) and P-values are indicated in the figures. In box and whiskers plots, horizontal lines of the box represent Q_3_, median and Q_1_, + represents the mean, and whiskers represent maximum and minimum values.

## Acknowledgements

We firstly would like to thank Dr. Isabel Fariñas, Dr. Anne Ferguson-Smith and Dr. Ángel Raya and their groups for technical support and discussion of the data. This work was supported by grants from Ministerio de Ciencia e Innovación/AEI (PID2019-110045GB-I00, PID2022-142734OB-I00 and EUR2023-143479), Generalitat Valenciana (AICO/2020/367) and Fundación BBVA to SRF. LLC (PRE2020-094137) and JDM (PRE2022-000680) were funded by the Spanish Formación de Personal Investigador (FPI) fellowship program EJV was funded by the Spanish Formación de Profesorado Universitario (FPU) fellowship program (FPU20/00795). ALU was funded by the Generalitat Valenciana fellowship program (ACIF/2016/381). Open Access funding was provided by the Ministerio de Ciencia e Innovación.

## Author Contribution

LLC, EJV, ALU, RML, JDM and ALP carried out most of the experiments. LLC performed gene expression and statistical analysis. EJV performed DNA methylation assays, helped by MI. ALU, RML and JDM performed NSC reprogramming. ALP contributed to develop the reprogramming protocol. JP performed the bioinformatic analysis of transcriptome and methylome data. EJR helped with MeDIP-seq bioinformatic analysis. SRF initiated, designed and led the study, and wrote the manuscript. All authors contributed to experimental design, data analysis, discussion and writing of the paper.

## Data availability

All relevant data can be found within the article and its supplementary information. Both methylation and expression data supporting the findings of this study have been deposited in Gene Expression Omnibus (GEO) with the accession numbers GSE282749 and GSE282748 respectively. RNA-seq raw counts and scaled MeDIP-seq signal tracks are provided as processed information in the GEO accession. A detailed list of tools used in the analysis of the present study can be found in Supplementary Table S9. Code developed to support current study can be provided upon request.

## References

Aasen T, Raya A, Barrero MJ, Garreta E, Consiglio A, Gonzalez F, Vassena R, Bilić J, Pekarik V, Tiscornia G, et al (2008) Efficient and rapid generation of induced pluripotent stem cells from human keratinocytes. Nat Biotechnol 26: 1276–1284

Alexander KA, Wang X, Shibata M, Clark AG & García-García MJ (2015) TRIM28 Controls Genomic Imprinting through Distinct Mechanisms during and after Early Genome-wide Reprogramming. Cell Rep 13: 1194–1205

Amemiya HM, Kundaje A & Boyle AP (2019) The ENCODE Blacklist: Identification of Problematic Regions of the Genome. Sci Rep 9: 9354

Apostolou E, Ferrari F, Walsh RM, Bar-Nur O, Stadtfeld M, Cheloufi S, Stuart HT, Polo JM, Ohsumi TK, Borowsky ML, et al (2013) Genome-wide Chromatin Interactions of the Nanog Locus in Pluripotency, Differentiation, and Reprogramming. Cell Stem Cell 12: 699–712

Arez M, Eckersley-Maslin M, Klobučar T, Lopes J von G, Krueger F, Mupo A, Raposo AC, Oxley D, Mancino S, Gendrel A-V, et al (2022) Imprinting fidelity in mouse iPSCs depends on sex of donor cell and medium formulation. Nat Commun 13: 5432

Barlow DP & Bartolomei MS (2014) Genomic imprinting in mammals. Cold Spring Harb Perspect Biol 6: a018382–a018382

Bartolomei MS & Ferguson-Smith AC (2011) Mammalian Genomic Imprinting. Cold Spring Harb Perspect Biol 3: a002592

Belenguer G, Domingo-Muelas A, Ferrón SR, Morante-Redolat JM & Fariñas I (2016) Isolation, culture and analysis of adult subependymal neural stem cells. Differentiation 91: 28–41

Benetatos L, Vartholomatos G & Hatzimichael E (2014) DLK1-DIO3 imprinted cluster in induced pluripotency: landscape in the mist. Cell Mol Life Sci 71: 4421–4430

Edwards CA & Ferguson-Smith AC (2007) Mechanisms regulating imprinted genes in clusters. Curr Opin Cell Biol 19: 281–289

Eminli S, Utikal J, Arnold K, Jaenisch R & Hochedlinger K (2008) Reprogramming of Neural Progenitor Cells into Induced Pluripotent Stem Cells in the Absence of Exogenous Sox2 Expression. STEM CELLS 26: 2467–2474

Ferguson-Smith AC (2011) Genomic imprinting: the emergence of an epigenetic paradigm. Nat Rev Genet 12: 565–575

Ferrón SR, Charalambous M, Radford E, McEwen K, Wildner H, Hind E, Morante-Redolat JM, Laborda J, Guillemot F, Bauer SR, et al (2011) Postnatal loss of Dlk1 imprinting in stem cells and niche-astrocytes regulates neurogenesis. Nature 475: 381–385

Frankish A, Diekhans M, Jungreis I, Lagarde J, Loveland JE, Mudge JM, Sisu C, Wright JC, Armstrong J, Barnes I, et al (2020) GENCODE 2021. Nucleic Acids Res 49: D916–D923

Gu Z, Eils R & Schlesner M (2016) Complex heatmaps reveal patterns and correlations in multidimensional genomic data. Bioinformatics 32: 2847–2849

Hackett JA, Sengupta R, Zylicz JJ, Murakami K, Lee C, Down TA & Surani MA (2013) Germline DNA Demethylation Dynamics and Imprint Erasure Through 5-Hydroxymethylcytosine. Science 339: 448–452

Hanna J, Carey BW & Jaenisch R (2008) Reprogramming of Somatic Cell Identity. Cold Spring Harb Symp Quant Biol 73: 147–155

Hashimoto H, Liu Y, Upadhyay AK, Chang Y, Howerton SB, Vertino PM, Zhang X & Cheng X (2012) Recognition and potential mechanisms for replication and erasure of cytosine hydroxymethylation. Nucleic Acids Res 40: 4841–4849

Hill PWS, Amouroux R & Hajkova P (2014) DNA demethylation, Tet proteins and 5-hydroxymethylcytosine in epigenetic reprogramming: An emerging complex story. Genomics 104: 324–333

Hochedlinger K & Jaenisch R (2015) Induced Pluripotency and Epigenetic Reprogramming. Cold Spring Harb Perspect Biol 7: a019448

Höpfl G, Gassmann M & Desbaillets I (2004) Differentiating embryonic stem cells into embryoid bodies. Methods Mol Biol (Clifton, NJ) 254: 79–98

Hotta A & Ellis J (2008) Retroviral vector silencing during iPS cell induction: An epigenetic beacon that signals distinct pluripotent states. J Cell Biochem 105: 940–948

Ito S, Shen L, Dai Q, Wu SC, Collins LB, Swenberg JA, He C & Zhang Y (2011) Tet Proteins Can Convert 5-Methylcytosine to 5-Formylcytosine and 5-Carboxylcytosine. Science 333: 1300–1303

Janiszewski A, Talon I, Chappell J, Collombet S, Song J, Geest ND, To SK, Bervoets G, Marin-Bejar O, Provenzano C, et al (2019) Dynamic reversal of random X-Chromosome inactivation during iPSC reprogramming. Genome Res 29: 1659–1672

Kim JB, Zaehres H, Wu G, Gentile L, Ko K, Sebastiano V, Araúzo-Bravo MJ, Ruau D, Han DW, Zenke M, et al (2008a) Pluripotent stem cells induced from adult neural stem cells by reprogramming with two factors. Nature 454: 646–650

Kim JB, Zaehres H, Wu G, Gentile L, Ko K, Sebastiano V, Araúzo-Bravo MJ, Ruau D, Han DW, Zenke M, et al (2008b) Pluripotent stem cells induced from adult neural stem cells by reprogramming with two factors. Nature 454: 646–650

Kim MJ, Choi HW, Jang HJ, Chung HM, Arauzo-Bravo MJ, Schöler HR & Do JT (2013) Conversion of genomic imprinting by reprogramming and redifferentiation. J Cell Sci 126: 2516–2524

Kurtenbach S & Harbour JW (2019) SparK: A Publication-quality NGS Visualization Tool. bioRxiv: 845529

Lassi G & Tucci V (2019) Genomic imprinting and the control of sleep in mammals. Curr Opin Behav Sci 25: 77–82

Lee HJ, Choi NY, Lee S-W, Ko K, Hwang TS, Han DW, Lim J, Schöler HR & Ko K (2016) Epigenetic alteration of imprinted genes during neural differentiation of germline-derived pluripotent stem cells. Epigenetics 11: 177–183

Lee HJ, Hore TA & Reik W (2014) Reprogramming the Methylome: Erasing Memory and Creating Diversity. Cell Stem Cell 14: 710–719

Lee S, Cook D & Lawrence M (2019a) plyranges: a grammar of genomic data transformation. Genome Biol 20: 4

Lee S, Cook D & Lawrence M (2019b) plyranges: a grammar of genomic data transformation. Genome Biol 20: 4

Li W, Zhao X, Wan H, Zhang Y, Liu L, Lv Z, Wang X-J, Wang L & Zhou Q (2011) iPS cells generated without c-Myc have active Dlk1-Dio3 region and are capable of producing full-term mice through tetraploid complementation. Cell Res 21: 550–553

Li X, Li MJ, Yang Y & Bai Y (2019) Effects of reprogramming on genomic imprinting and the application of pluripotent stem cells. Stem Cell Res 41: 101655

Lian H, Li W-B & Jin W-L (2016) The emerging insights into catalytic or non-catalytic roles of TET proteins in tumors and neural development. Oncotarget 7: 64512–64525

Lienhard M, Grimm C, Morkel M, Herwig R & Chavez L (2014) MEDIPS: genome-wide differential coverage analysis of sequencing data derived from DNA enrichment experiments. Bioinformatics 30: 284–286

Liu L, Luo G-Z, Yang W, Zhao X, Zheng Q, Lv Z, Li W, Wu H-J, Wang L, Wang X-J, et al (2010) Activation of the Imprinted Dlk1-Dio3 Region Correlates with Pluripotency Levels of Mouse Stem Cells. J Biol Chem 285: 19483–19490

Love MI, Huber W & Anders S (2014) Moderated estimation of fold change and dispersion for RNA-seq data with DESeq2. Genome Biol 15: 550

Love MI, Soneson C, Hickey PF, Johnson LK, Pierce NT, Shepherd L, Morgan M & Patro R (2020) Tximeta: Reference sequence checksums for provenance identification in RNA-seq. PLoS Comput Biol 16: e1007664

Messerschmidt D (2012) Should I stay or should I go: Protection and maintenance of DNA methylation at imprinted genes. Epigenetics 7: 969–975

Mikkelsen TS, Hanna J, Zhang X, Ku M, Wernig M, Schorderet P, Bernstein BE, Jaenisch R, Lander ES & Meissner A (2008) Dissecting direct reprogramming through integrative genomic analysis. Nature 454: 49–55

Montalbán-Loro R, Lassi G, Lozano-Ureña A, Perez-Villalba A, Jiménez-Villalba E, Charalambous M, Vallortigara G, Horner AE, Saksida LM, Bussey TJ, et al (2021) Dlk1 dosage regulates hippocampal neurogenesis and cognition. Proc Natl Acad Sci 118: e2015505118

Montalbán-Loro R, Lozano-Ureña A, Ito M, Krueger C, Reik W, Ferguson-Smith AC & Ferrón SR (2019) TET3 prevents terminal differentiation of adult NSCs by a non-catalytic action at Snrpn. Nat Commun 10: 1726

Nazor KL, Altun G, Lynch C, Tran H, Harness JV, Slavin I, Garitaonandia I, Müller F-J, Wang Y-C, Boscolo FS, et al (2012) Recurrent Variations in DNA Methylation in Human Pluripotent Stem Cells and Their Differentiated Derivatives. Cell Stem Cell 10: 620–634

Parry A, Rulands S & Reik W (2021) Active turnover of DNA methylation during cell fate decisions. Nat Rev Genet 22: 59–66

Patro R, Duggal G, Love MI, Irizarry RA & Kingsford C (2017) Salmon provides fast and bias-aware quantification of transcript expression. Nat Methods 14: 417–419

Perez JD, Rubinstein ND & Dulac C (2016) New Perspectives on Genomic Imprinting, an Essential and Multifaceted Mode of Epigenetic Control in the Developing and Adult Brain. Annu Rev Neurosci 39: 347–84

Perrera V & Martello G (2019) How Does Reprogramming to Pluripotency Affect Genomic Imprinting? Front Cell Dev Biol 7: 76

Pham A, Selenou C, Giabicani E, Fontaine V, Marteau S, Brioude F, David L, Mitanchez D, Sobrier ML & Netchine I (2022) Maintenance of methylation profile in imprinting control regions in human induced pluripotent stem cells. Clin Epigenetics 14: 190

Prokhorova TA, Harkness LM, Frandsen U, Ditzel N, Schrder HD, Burns JS & Kassem M (2009) Teratoma Formation by Human Embryonic Stem Cells Is Site Dependent and Enhanced by the Presence of Matrigel. Stem Cells Dev 18: 47–54

Ramírez F, Dündar F, Diehl S, Grüning BA & Manke T (2014) deepTools: a flexible platform for exploring deep-sequencing data. Nucleic Acids Res 42: W187–W191

Robinson MD, McCarthy DJ & Smyth GK (2009) edgeR: a Bioconductor package for differential expression analysis of digital gene expression data. Bioinformatics 26: 139–140

SanMiguel JM & Bartolomei MS (2018) DNA methylation dynamics of genomic imprinting in mouse development†. Biol Reprod 99: 252–262

Santini L, Halbritter F, Titz-Teixeira F, Suzuki T, Asami M, Ma X, Ramesmayer J, Lackner A, Warr N, Pauler F, et al (2021) Genomic imprinting in mouse blastocysts is predominantly associated with H3K27me3. Nat Commun 12: 3804

Sardina JL, Collombet S, Tian TV, Gómez A, Stefano BD, Berenguer C, Brumbaugh J, Stadhouders R, Segura-Morales C, Gut M, et al (2018) Transcription Factors Drive Tet2-Mediated Enhancer Demethylation to Reprogram Cell Fate. Cell Stem Cell 23: 727–741.e9

Silva J, Barrandon O, Nichols J, Kawaguchi J, Theunissen TW & Smith A (2008) Promotion of Reprogramming to Ground State Pluripotency by Signal Inhibition. PLoS Biol 6: e253

Smallwood SA & Kelsey G (2012) De novo DNA methylation: a germ cell perspective. Trends Genet 28: 33–42

Spelke DP, Ortmann D, Khademhosseini A, Ferreira L & Karp JM (2010) Methods for embryoid body formation: the microwell approach. Methods Mol Biol (Clifton, NJ) 690: 151–62

Stadtfeld M, Apostolou E, Akutsu H, Fukuda A, Follett P, Natesan S, Kono T, Shioda T & Hochedlinger K (2010) Aberrant silencing of imprinted genes on chromosome 12qF1 in mouse induced pluripotent stem cells. Nature 465: 175–181

Stadtfeld M, Brennand K & Hochedlinger K (2008) Reprogramming of pancreatic beta cells into induced pluripotent stem cells. Curr Biol : CB 18: 890–4

Tahiliani M, Koh KP, Shen Y, Pastor WA, Bandukwala H, Brudno Y, Agarwal S, Iyer LM, Liu DR, Aravind L, et al (2009) Conversion of 5-Methylcytosine to 5-Hydroxymethylcytosine in Mammalian DNA by MLL Partner TET1. Science 324: 930–935

Taiwo O, Wilson GA, Morris T, Seisenberger S, Reik W, Pearce D, Beck S & Butcher LM (2012) Methylome analysis using MeDIP-seq with low DNA concentrations. Nat Protoc 7: 617–636

Takahashi K & Yamanaka S (2006) Induction of Pluripotent Stem Cells from Mouse Embryonic and Adult Fibroblast Cultures by Defined Factors. Cell 126: 663–676

Takahashi N, Gray D, Strogantsev R, Noon A, Delahaye C, Skarnes WC, Tate PH & Ferguson-Smith AC (2015) ZFP57 and the Targeted Maintenance of Postfertilization Genomic Imprints. Cold Spring Harb Symp Quant Biol 80: 177–187

Takikawa S, Ray C, Wang X, Shamis Y, Wu T-Y & Li X (2013) Genomic imprinting is variably lost during reprogramming of mouse iPS cells. Stem Cell Res 11: 861–873

Tucci V, Isles AR, Kelsey G, Ferguson-Smith AC, Group the EI, Tucci V, Bartolomei MS, Benvenisty N, Bourc’his D, Charalambous M, et al (2019) Genomic Imprinting and Physiological Processes in Mammals. Cell 176: 952–965

Vaz IM, Borgonovo T, Kasai-Brunswick TH, Santos DS dos, Mesquita FCP, Vasques JF, Gubert F, Rebelatto CLK, Senegaglia AC & Brofman PRS (2021) Chromosomal aberrations after induced pluripotent stem cells reprogramming. Genet Mol Biol 44: e20200147

Vu H & Ernst J (2023) Universal chromatin state annotation of the mouse genome. Genome Biol 24: 153

Wickham H, Averick M, Bryan J, Chang W, McGowan L, François R, Grolemund G, Hayes A, Henry L, Hester J, et al (2019) Welcome to the Tidyverse. J Open Source Softw 4: 1686

Wilson GA, Dhami P, Feber A, Cortázar D, Suzuki Y, Schulz R, Schär P & Beck S (2012) Resources for methylome analysis suitable for gene knockout studies of potential epigenome modifiers. GigaScience 1: 3

Wu T, Hu E, Xu S, Chen M, Guo P, Dai Z, Feng T, Zhou L, Tang W, Zhan L, et al (2021) clusterProfiler 4.0: A universal enrichment tool for interpreting omics data. Innov 2: 100141

Wu X & Zhang Y (2017) TET-mediated active DNA demethylation: mechanism, function and beyond. Nat Rev Genet 18: 517–534

Yagi M, Kabata M, Ukai T, Ohta S, Tanaka A, Shimada Y, Sugimoto M, Araki K, Okita K, Woltjen K, et al (2019) De Novo DNA Methylation at Imprinted Loci during Reprogramming into Naive and Primed Pluripotency. Stem Cell Rep 12: 1113–1128

Ying Q-L, Stavridis M, Griffiths D, Li M & Smith A (2003) Conversion of embryonic stem cells into neuroectodermal precursors in adherent monoculture. Nat Biotechnol 21: 183–186

Ying Q-L, Wray J, Nichols J, Batlle-Morera L, Doble B, Woodgett J, Cohen P & Smith A (2008) The ground state of embryonic stem cell self-renewal. Nature 453: 519–523

Yu G, Wang L-G & He Q-Y (2015) ChIPseeker: an R/Bioconductor package for ChIP peak annotation, comparison and visualization. Bioinformatics 31: 2382–2383

Zimmerman DL, Boddy CS & Schoenherr CS (2013) Oct4/Sox2 Binding Sites Contribute to Maintaining Hypomethylation of the Maternal Igf2/H19 Imprinting Control Region. PLoS ONE 8: e81962

Zou Z, Ohta T & Oki S (2024) ChIP-Atlas 3.0: a data-mining suite to explore chromosome architecture together with large-scale regulome data. Nucleic Acids Res 52: W45–W53

